# Persistent Japanese encephalitis virus infection in fetuses of an amplifying pig host affects virus genetic heterogeneity and drives transcriptional footprints in offspring

**DOI:** 10.64898/2026.09.05.749582

**Authors:** Prince Pal Singh, Ivan Trus, Daniel Udenze, Brian Cox, Uladzimir Karniychuk

## Abstract

Japanese encephalitis virus (JEV) poses a global threat to public health. This study demonstrates that during fetal infection in natural amplifying hosts—pigs, the *in utero* environment is conducive to the emergence of new intra-host JEV variants. Transplacental and fetal infection in pigs with JEV resulted in the emergence of virus variants that at least partially resembled variants reported in the field samples from JEV endemic regions. The JEV genetic heterogeneity acquired during fetal infection persisted in expelled fetal membranes and offspring. Like in fetuses, JEV mutations identified in afterbirth fetal membranes and offspring were previously reported in the field. Several selected fetal- and offspring-specific mutations that emerged during persistent *in utero* infection were stable during passage in cell culture. This comparative analysis with field-reported mutations, together with the stability of emerged mutations during serial passages, may suggest the potential for onward transmission of *in utero*-emerged JEV variants from infected fetal membranes or offspring. Exposure to JEV during fetal life also induced transcriptional footprints in blood cells and tonsils of live offspring, including alterations in pathways associated with epigenetic regulation and memory. Altogether, we generated new knowledge on JEV biology and pathogenesis in its natural amplifying hosts—pigs—during pregnancy. Our findings provide directions for future studies investigating pathways linking fetal infection to offspring and subsequent environmental dissemination, fetal infection as an additional source of JEV genetic diversification, and the mechanisms underlying molecular footprints and their long-term sequelae in JEV-affected offspring, including increased offspring susceptibility to zoonotic pathogens.

**Author summary:** Japanese encephalitis virus (JEV) is the leading cause of encephalitis in humans in the Asia-Pacific region and a significant threat to food security due to its circulation in pig farms in endemic areas. Pigs are the amplifying host for JEV, and proximity to pig farms is a major risk for human infection. The JEV pathogenesis in pregnant pigs remains poorly understood. This is an important knowledge gap, as transplacental infection and persistence in fetuses during prolonged porcine gestation may constitute a distinct pathogenic, evolutionary, and transmission mechanisms for JEV. Here, we profiled JEV genetic heterogeneity during persistent infection in fetuses and offspring and identified the emergence of new viral variants. We also demonstrated that clinically healthy offspring born to JEV-exposed dams carry transcriptional footprints, indicating lasting effects of exposure during fetal life. Our findings open new directions for research on how fetal infection may contribute to JEV spread after birth, emergence of new viral variants, and increased susceptibility of affected piglets to other zoonotic pathogens.

## Introduction

Nearly half of the world’s population lives in territories where Japanese encephalitis virus (JEV) is endemic [1]. JEV is the leading cause of human encephalitis in the Asia-Pacific region [2, 3]. There is a concern that JEV can be introduced into North America given a large population of amplifying hosts—domestic and wild pigs; as well as of susceptible *Culex* mosquitoes [4–6]. JEV-infected piglets typically do not die but may develop fever and tremors [7–10]. While mortality in adult pigs is negligible, they can develop viremia and persistent infection [11, 12]. This combination of high survival and high susceptibility makes pigs efficient amplifying hosts, facilitating zoonotic spillover of JEV to humans. Indeed, proximity to pig farms is a major risk for JEV infections in humans [13, 14]. There is increasing evidence showing that JEV persistence in pig herds has high complexity and plays an important role in its biology. Pig herds can sustain JEV between mosquito seasons in endemic regions [15], likely through pig-to-pig contact transmission [8, 12]. However, the role of specific age groups of pigs and physiological states, such as pregnancy, is also not well understood.

The pathogenesis and role of pregnant pigs in JEV transmission are understudied. Here, we capitalized on an existing inventory of JEV-infected fetal samples from our previous study where fetuses were sampled before birth [12] and conducted an additional experiment in which pregnant pigs infected with JEV at mid-gestation delivered piglets after more than two months of persistent fetal infection. This experimental design provided a unique opportunity to characterize and compare whole-genome JEV heterogeneity in developing fetuses and newborn offspring. Furthermore, we assessed whether JEV infection during fetal life induces transcriptional footprints in blood cells and immune organs of surviving piglets, potentially increasing their susceptibility to other infections after birth.

## Materials and Methods

### Ethics statement

For pig experiments, we followed the Canadian Council on Animal Care guidelines and Animal Use Protocols #20200106 and #20230070 approved by the University of Saskatchewan’s Animal Research Ethics Board. All efforts were made to minimize animal suffering. Pigs were euthanized with an anesthetic overdose followed by exsanguination. BSL3 JEV work was approved by the University of Saskatchewan Biosafety Permit #I-IVC-10. The JEV studies were also approved by The Ohio State University Institutional Biosafety Committee Protocols #2023R00000075 and #2022R00000104.

For work involving human umbilical cord blood samples, we followed the University of Saskatchewan Human Research Ethics Policy, with approval from Biomedical Research Ethics Board (Bio-REB) #16-135. Pregnant women recruited (27^th^ August 2020–25^th^ November 2020) into the study (see Supplementary Materials and Methods, **S1 File**) for umbilical cord blood donation provided written informed consent.

### Viruses and cells

The JEV Nakayama strain [GenBank: #EF571853.1] stock was initially produced at the World Reference Center for Emerging Viruses and Arboviruses at the University of Texas Medical Branch at Galveston and transferred to our laboratory through the Public Health Agency of Canada. We inoculated Vero E6 cells and harvested media at 9 days after inoculation to produce the working stock. Culture media containing JEV were centrifuged (12,000g, 20 min, +4°C); the supernatant was collected, aliquoted, and frozen at −80°C. BHK-21 (ATCC #CCL-10) and Vero E6 (ATCC #CRL-1586) cells were cultured in DMEM (Fisher, MA, USA; #11-965-118) supplemented with 3% FBS (Fisher, MA, USA; #A5256801), 1× Penicillin-streptomycin (Fisher, MA, USA; #15140122), and 2.67 mM Sodium bicarbonate (Fisher, MA, USA; #25080094). Cells and JEV stock were mycoplasma-free as confirmed by the LookOut Mycoplasma PCR Detection Kit (Millipore Sigma, MA, USA; #MP0035).

### Pig experiments

Landrace-cross pigs were purchased from the university high-health status herd free from clinical signs of porcine reproductive and respiratory virus (PRRSV), porcine parvovirus (PPV), congenital porcine circovirus 2 (PCV2), and porcine circovirus 3 (PCV3), which can cause fetal infection in pigs. Accordingly, maternal and selected fetal and piglet samples were negative for PRRSV, PPV, PCV2, and PCV3 in virus-specific PCR assays [16, 17]. Before delivering to containment, pigs were synchronized and bred with semen from a single donor to reduce biological variability. All pigs used for fetal or offspring studies were housed simultaneously under identical conditions.

The details of the ***fetal experiment*** were previously published [12]. Briefly, insemination was scheduled to ensure that at the time of pig inoculation with JEV, three stages of pregnancy (the duration of pregnancy in pigs is 114 days) were represented: two pigs at early pregnancy (inoculation at 30 days of pregnancy), two pigs at mid-pregnancy (54 days), and two pigs at late pregnancy (86 days). Two non-pregnant adult pigs were also included. The pregnancy was confirmed with ultrasound imaging, and all pigs were delivered to Vaccine and Infectious Disease Organization, University of Saskatchewan BSL3 containment facility. Animals were housed in identical rooms in individual pens with no contact with each other. After seven days of acclimatization in containment, all pigs were sedated and inoculated with 10^7^ TCID_50_ of JEV intradermally (ear skin, 1 ml) + intravenous (ear vein, 1ml). This inoculation dose and routes were the same as previously used for JEV inoculation in young piglets [18, 19]; in addition to modeling JEV infection in pig fetuses and offspring, this approach enables direct comparison with prior studies in young piglets. Clinical signs, including appetite, activity, and rectal temperature, were recorded before and after JEV inoculation [12]. In the fetal study, one non-pregnant pig showed elevated body temperature 39.3–39.4°C (the baseline temperature was 38.4–38.5°C) and decreased appetite at 3–4 days after JEV inoculation. Other animals did not show clinical signs.

We collected blood from the jugular vein with BD Vacutainer™ Plastic Blood Collection EDTA tubes, as well as nasal and vaginal swabs. Samples were collected before JEV inoculation and at 1–7, 14, 21, and 28 days after virus inoculation. After blood centrifugation (2,000g, 20 min, +4 °C), plasma was aliquoted and frozen at −80°C. For nasal and vaginal swabs, swabs were inserted to the nose or vagina and rotated to obtain secretions. Afterward swabs were placed into tubes containing 500 µl sterile media, the handle was broken one centimeter from the top of the swab, and the tube was stored at −80°C.

Pigs were euthanized and sampled 28 days after JEV inoculation. In six pregnant pigs, uteri with fetuses were removed to sample each fetus with individual sterile instruments. Umbilical cord blood was aspirated from each fetus with sterile syringes and needles, centrifuged, and plasma was aliquoted and frozen at −80°C.

In the initial fetal study, maternal, transplacental, and fetal infections were confirmed and characterized. This included JEV RNA loads in multiple maternal tissues; productive transplacental infection evidenced by high JEV RNA levels in fetal plasma; the presence of infectious JEV in fetal plasma; JEV protein expression in fetal brains; and severe developmental pathology in a subset of infected fetuses [12]. JEV-positive maternal blood, vaginal and nasal swabs, as well as fetal blood were used for NGS in the present study.

For the ***offspring experiment*** pregnant and non-pregnant pigs bred in the same university high-health status herd as for fetal study were used. The pregnancy was confirmed with ultrasound, and all pigs were delivered to Vaccine and Infectious Disease Organization, University of Saskatchewan BSL-3 containment facility. Insemination was scheduled to obtain seven pregnant pigs at mid-pregnancy (50 days of gestation). Animals were housed in identical rooms in individual pens with no contact with each other. After seven days of acclimatization in containment, six pigs were sedated and inoculated with 10^7^ TCID_50_ of JEV as in the fetal study. One control pig was mock-inoculated with the vehicle using the same procedure. Clinical signs, including appetite, activity, and rectal temperature, were recorded before and after JEV inoculation [12], and adult pigs did not show clinical signs.

We collected blood from the jugular vein of adult pigs with BD Vacutainer™ Plastic Blood Collection EDTA tubes. Based on detectable JEV viremia defined in the fetal study [12], blood samples were collected before JEV inoculation and at 1, 2, and 3 days after virus inoculation. After blood centrifugation (2,000g, 20 min, +4°C), plasma was aliquoted and frozen at −80°C.

All pregnant pigs were monitored until delivery at 115-116 days of gestation. Piglets were sampled at birth (dead piglets) or at 3–4 days of age (live piglets; all in Control and Severe subgroups, or 12 selected piglets in the Mild subgroup) for JEV quantification and NGS analysis. Specifically, samples included whole blood (EDTA tubes for plasma and Tempus Blood RNA Tubes for whole blood), amniotic membrane, placenta, brain, nasal swabs, and tonsils.

### Statistical analysis

We used GraphPad PRISM 8 software and R v4.2 to analyze and visualize data. The difference with *P* < 0.05 was considered statistically significant. All data were expressed as individual values with mean ± standard deviation (M ± SD). The number of CFU-GM, GMPs and IL-1*β* levels were compared between different cellular conditions with Kruskal–Wallis H-test. Statistical methods used for JEV nucleotide diversity and RNA-seq analyses are in **S1 File**.

Details of RNA extraction, RT-qPCR, JEV negative-strand-RNA-specific RT-PCR; classical non-targeted and PrimalSeq NGS protocols and Sanger sequencing; Illumina data processing and variant calling; JEV nucleotide diversity analysis; reverse genetics to rescue fetal and offspring-specific JEV variants; JEV serial passaging for acquired mutation stability assays; RNA-seq and gene expression analysis; hematopoietic stem/progenitor cell (HSPC) experiments; and whole-genome DNA methylation array and data processing are provided in **S1 File** (Supplementary Materials and Methods).

## Results

### JEV acquires genetic heterogeneity during fetal infection

The JEV Nakayama strain, which we used for injection of pregnant pigs, was isolated from the cerebrospinal fluid of a patient in 1935. The reference sequence of the full Nakayama genome is available in NCBI [GenBank: EF571853.1] and dated by May 2007. Thus, we questioned whether the available whole-genome JEV Nakayama NCBI from 2007 sequence can serve as a suitable reference for JEV evolutionary studies. We used well-established techniques in our laboratory, previously adapted for whole-genome NGS of West Nile virus and Zika virus [12, 16, 20–22]. First, we extracted the RNA from the JEV Nakayama stock, which was used for inoculation of pregnant pigs, and applied the NGS PrimalSeq protocol, the same approach that we used for NGS in pig tissues (see below and Supplementary Materials and Methods, **S1 File**). Upon aligning PrimalSeq-derived JEV stock NGS reads with the historical 2007 JEV Nakayama NCBI sequence [GenBank: EF571853.1], we identified 69 SNVs (**Fig 1** and **S1 Table**). 59 SNVs were dominant with almost 100% of frequency. The other 10 SNVs showed frequencies between 4% and 72% (**Fig 1B**). Second, to validate whether identified SNVs are either true or false positives, we generated a consensus whole-genome JEV sequence directly from the virus stock (which was used for inoculation of pigs) using untargeted NGS (Supplementary Materials and Methods, **S1 File**). In contrast to samples from JEV infected pigs, the virus stock derived from the cell culture had high viral loads, permitting the use of the untargeted NGS protocol without advanced PrimalSeq PCR pre-amplification. This approach resulted in an in-house whole-genome JEV Nakayama reference (**S2 File**). When we compared PrimalSeq NGS JEV stock reads with the in-house whole-genome JEV Nakayama reference (**S2 File**), we identified only 10 SNVs (**Fig 1** and **S1 Table**). Two SNVs had 15% and 28% frequency; frequencies in the remaining 8 SNVs ranged from 4% to 10%. These PrimalSeq 10 SNVs are most probably a result of how software generates the in-house whole-genome JEV Nakayama consensus reference sequence (**S2 File**). The in-house whole-genome JEV Nakayama consensus reference sequence is based on a Unipro UGENE algorithm which uses ≥50% major-allele threshold [23] (Supplementary Materials and Methods, **S1 File**); bases below this cutoff are ignored during consensus construction. Thus, the NCBI-derived JEV reference sequence [GenBank: EF571853.1] is not well-suited for accurate analysis of intra-host JEV evolution, can lead to false discovery of SNVs in experimental tissues, and thereby requiring the generation of an in-house reference stock sequence (**S2 File**).

**Fig 1.**
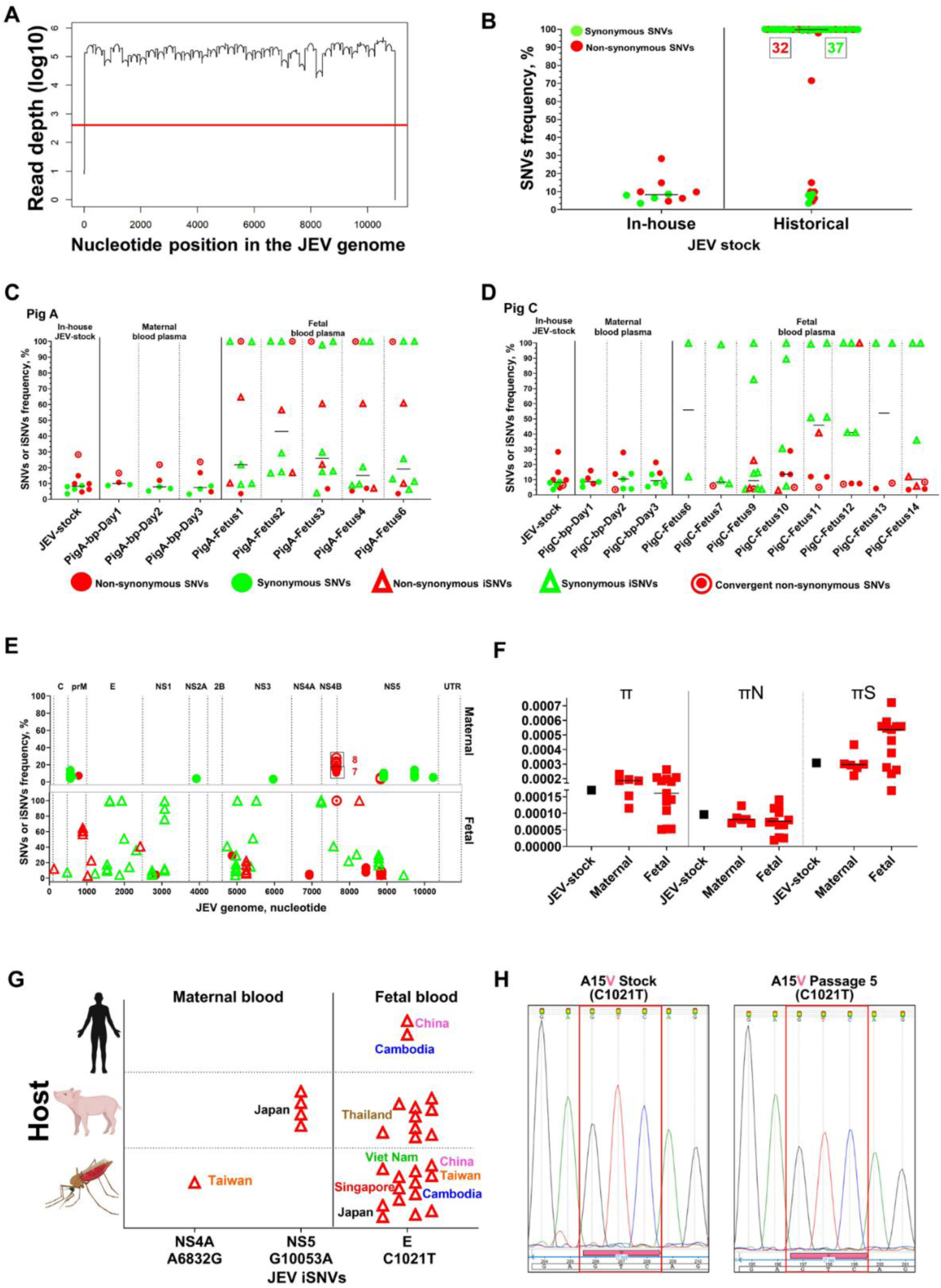
JEV evolution in maternal and fetal compartments. **(A)** JEV stock genome coverage and depth obtained with the PrimalSeq NGS protocol. The red horizontal line shows 400 nucleotide NGS depth used as a threshold for accurate SNV identification. **(B)** Number and frequency of SNVs in the JEV stock after alignment to the in-house (**S2 File**) or historical [GenBank: #EF571853.1] reference JEV Nakayama sequence. Green or red numbers represent the total number of synonymous or non-synonymous SNVs, respectively. **(C)** Data from pig #A inoculated with JEV at early pregnancy (30 days), and **(D)** pig #C inoculated with JEV at mid-pregnancy (54 days). Both pigs were sampled at 28 days after inoculation; the total duration of pregnancy in pigs is 114–116 days. **SNVs:** Single nucleotide variants. **iSNVs:** *de novo* emerged intra-host SNVs. Solid horizontal line - median values. Details of all SNVs and iSNVs are in **S1 Table**. (**E**) SNV distribution pattern in the JEV genome identified in the maternal and fetal blood. C: JEV genomic region encoding capsid protein; prM: precursor membrane protein; E: envelope protein; NS1: nonstructural protein 1; NS2A: nonstructural protein 2A; NS2B: nonstructural protein 2B; NS3: nonstructural protein 3; NS4A: nonstructural protein 4A; NS4B: nonstructural protein 4B; NS5: nonstructural protein 5. UTR: untranslated region. Raw data are shown in **S1 Table**. (**F**) Nucleotide diversity metrics. X-axis represents sample type. Black—JEV stock; red—blood plasma. Y-axis: Mean pairwise difference for π (mean number of pairwise differences per site across the whole genome), πN (mean number of pairwise non-synonymous differences per non-synonymous site across the genome) and πS (mean number of pairwise synonymous differences per synonymous site across the genome). Solid horizontal lines represent median values. Raw data are shown in **S1 Table**. (**G**) Comparative experimental and field JEV evolution; **S1** and **S4 Tables**. Maternal and fetal *de novo* emerged **non-synonymous iSNVs** identified in this study and previously reported in field studies (NCBI database; see Supplementary Materials and Methods and **S4 Table**). Red empty triangles represent previously reported field occurrences of iSNVs identified in this study. X-axis represents specific iSNV and their position within the JEV genome. E: Envelope protein. NS4A and NS5: Nonstructural proteins 4A and NS5. (**H**) Sanger sequencing. Fetal *de novo* emerged non-synonymous iSNV C1021T (amino acid substitution A15V) identified in this study and previously reported in the field studies (**S4 Table**). Introduced A15V substitution showed stability after 5 passages in Vero cells. Nucleotide peaks highlighted in red squares encode the introduced A15V substitution. Raw FASTQ NGS files are deposited in BioProject: #PRJNA1467537. Portions of the figure are created in BioRender: Karniychuk, V. (2026) https://BioRender.com/xw9t9tk.

Maternal samples from six pregnant (#A-#F) and two non-pregnant adult pigs (#G, #H) infected with JEV were obtained from our previous study [12] (**S1 File**). Viral loads in maternal blood, vaginal, and nasal swabs [12] which were used for NGS in this study are summarized in **S2 Table**. We identified and analyzed SNVs and *de novo* emerged intra-host SNVs (iSNVs) with an open-source software package iVar. For SNVs, we compared NGS reads from the virus stock and pig samples to the in-house consensus reference JEV sequence (**S2 File**). Synonymous and non-synonymous mutations which were not present in the reference JEV sequence were considered as SNVs. Then, we compared the genomic positions and frequencies of SNVs between sequences from the JEV virus stock used for pig inoculation and JEV sequences from pig tissues; tissue-specific SNVs that were not present in initial JEV stock were considered as *de novo* emerged iSNVs.

Library construction and NGS were successful for both blood and nasal swabs in 2 out of 8 adult animals (pigs #C and #E; **S1 Fig**), and for both blood and vaginal swabs in 6 out of 8 adult pigs (pigs #A, #B, #C, #D, #F, and #H; **S1 Fig**). The percentage of *de novo* emerged iSNVs (triangles in **S1 Fig**) calculated from the total number of sample-specific mutations was considerably higher in nasal swabs and vaginal swabs than in blood in both adult pigs (**S1 Fig**).

Two pregnant pigs (#A and #C) from our previous study (**S2 Table**) [12] developed transplacental and fetal infections on sampling day 28 after maternal JEV inoculation (pregnant pigs were inoculated on day 30 or 54 of pregnancy; the duration of pregnancy in pigs is 114 days). In addition to the detection of JEV RNA in multiple maternal tissues (**S2 Table**), productive transplacental infection was confirmed by high JEV RNA loads in fetal blood plasma (**S2 Table**), the presence of infectious JEV in fetal plasma, JEV protein expression in fetal brains, and severe developmental pathology in a subset of infected fetuses [12]. Viral loads in fetal blood samples which were used for NGS in this study are summarized in **S2 Table**.

For pig #A, the percentage of fetal SNVs (solid colored dots in **Fig 1C**) calculated from the total number of sample-specific mutations was considerably lower than the percentage of fetal iSNVs (solid colored triangles in **Fig 1C**): fetus #1—SNVs 22% versus iSNVs 78%, fetus #2—12% versus 88%, fetus #3—20% versus 80%, fetus #4—30% versus 70%, and fetus #6—20% versus 80%. For pig #C, the percentage of fetal SNVs (solid colored dots in **Fig 1D**) calculated from the total number of sample-specific mutations was also considerably lower than the percentage of fetal iSNVs (solid colored triangles in **Fig 1D**): fetus #6—SNVs 0% versus iSNVs 100%, fetus #7—25% versus 75%, fetus #9—10% versus 90%, fetus #10—44% versus 56%, fetus #11—37% versus 63%, fetus #12—37% versus 63%, fetus #13—50% versus 50%, and fetus #14—37% versus 63%.

For pig #A, fetal iSNVs were located in the JEV genomic regions encoding prM, E, NS1, NS3, NS4B and NS5 proteins (**S2 Fig**). For pig #C, fetal iSNVs were located in the JEV genomic regions encoding prM, E, NS1, NS3, and NS5 proteins (**S2 Fig**).

To quantify the degree of convergent evolution in the JEV genome, any iSNV that emerged in more than one fetus was defined as convergent, and the total convergence was quantified as the percentage of convergent iSNVs relative to the total number of iSNVs. In fetuses belonging to pig #A, the degree of convergent evolution was 22%, synonymous iSNVs—16%, and non-synonymous iSNVs—5% (**S2 Fig** and **S1 Table**). In fetuses from pig #C, the degree of convergent evolution was 8.33% and all convergent iSNVs were synonymous (**S2 Fig** and **S1 Table**). Interestingly, a convergent non-synonymous SNV (A7656G corresponding to N131D amino acid substitution) was observed within NS4B region of the JEV genome in both maternal blood and blood from all the fetuses belonging to pig #A (**S2 Fig** and **S1 Table**). The frequency of this specific non-synonymous SNV was similar in inoculation JEV stock (28%) and maternal blood (average 21%) (**Fig 1C** and **S2 Fig**). However, it showed a considerable increase, rising from 21% in maternal blood to 100% in the blood plasma of all five fetuses (**Fig 1C**, **S2 Fig, S1 Table**). This increase may suggest distinct maternal and fetal evolutionary forces and an evolutionary placental bottleneck during transplacental transmission, as the other 10 JEV stock-specific and 13 maternal blood-specific SNVs (**Fig 1C**) were not present in fetal samples.

The low nucleotide diversity (π) across maternal and fetal samples (**Fig 1F**) suggests that overall viral populations remain genetically constrained within both biological compartments. Synonymous nucleotide diversity (πS) also exceeded non-synonymous nucleotide diversity (πN) in both compartments (**Fig 1F**), consistent with purifying selection [24]. However, despite the overall low diversity, the consistent emergence of compartment-specific iSNVs in fetuses (**Fig 1C**–**1E**) indicates ongoing localized virus evolution and the emergence or enrichment of potentially biologically meaningful JEV variants. To partially test this, we compared iSNVs that emerged during JEV infection in pregnant pigs and their fetuses (**S1 Table**) with JEV sequences available in the NCBI nucleotide database. For maternal samples, two *de novo* emerged non-synonymous iSNVs (A6832G corresponding to E123G amino acid substitution; and G10053A corresponding to V793M amino acid substitution) were reported during 1998-2016 in JEV sequences from Taiwan and Japan and identified in field samples from mosquitoes and pig sera (**Fig 1G** and **S4 Table**). Interestingly, in fetal samples, a *de novo* non-synonymous iSNV C1021T (A15V amino acid substitution) has been reported in JEV sequences from multiple geographic regions and hosts, including humans, mosquitoes, and pigs (**Fig 1G**, **S4 Table**). We used reverse genetics to synthetically rescue a JEV variant containing the C1021T mutation corresponding to A15V amino acid substitution (the reason for selecting this mutation is provided in **S1 File**), passaged the virus five times in cell culture, and confirmed stability of the fetal-specific mutation by Sanger sequencing (**Fig 1H**); the parental JEV variant retained the wild-type sequence before and after passaging.

Altogether, the *in utero* and fetal environment is conducive to the emergence of new intra-host JEV variants represented by synonymous and non-synonymous iSNVs in fetal blood (**Fig 1C,D**). Also, the experimental infection of pig fetuses with JEV resulted in the emergence of virus variants that at least partially resembled variants reported in the field samples from different JEV endemic regions (**Fig 1** and **S4 Table**).

### The JEV genetic heterogeneity acquired during fetal infection persists in expelled fetal membranes, dead fetuses, and surviving offspring

Next, we tested whether the heterogeneous JEV population that emerged *in utero* (**Fig 1**) can be shed into the environment after birth with expelled fetal membranes and by piglets infected during fetal development. We injected pregnant pigs with JEV at 50 days of gestation. Pregnant pigs were observed until delivery at 115–116 days of gestation, and piglets were sampled at birth or 3–4 days afterward (**Fig 2A**) for JEV quantification and NGS analysis.

**Fig 2.**
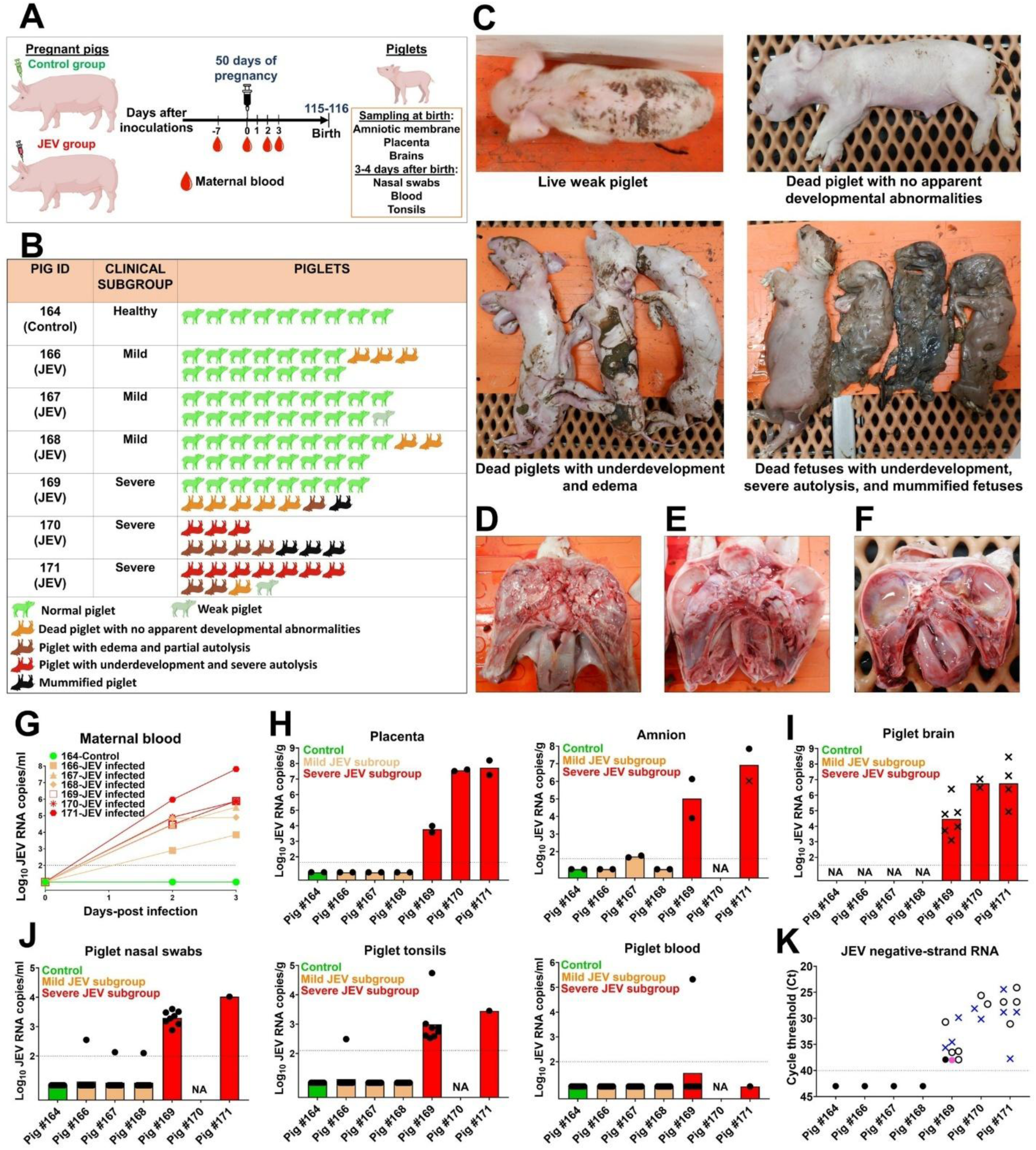
JEV infection in pregnant pigs and offspring. **(A)** Experimental design for offspring study. Seven pigs in the mid-pregnancy stage (50 days of pregnancy) were inoculated with media (control group) or JEV (JEV group). Experimental details are in Materials and Methods (**S1 File)**. **(B)** Clinical outcomes at birth. **(C)** Gross pathology in fetuses/piglets from JEV-infected pigs. **(D)** A skull containing the brain of a dead piglet with no apparent developmental abnormalities. **(E)** The skull of an underdeveloped dead piglet with edema showing partial brain autolysis. **(F)** The skull of an underdeveloped dead piglet showing full brain autolysis. **(G)** JEV RNA loads determined by virus-specific RT-qPCR in blood plasma of pregnant pigs at day 2 and 3 post JEV inoculation. Data for individual pregnant pigs; colors represent subgroups (also see **B**) identified based on pathology in piglets after birth: Control subgroup with healthy litter (green color), Mild clinical subgroup (brown) and Severe clinical subgroup (red). Different symbol shapes represent individual pregnant pigs. **(H-J)** JEV RNA loads determined by virus-specific RT-qPCR in samples from piglets; data are shown as mean (bar) and individual sample values (shapes). X-axis represents pregnant pig IDs and symbol shape represents individual liveborn (dots) or stillborn (crosses) piglets. Y-axis represents log_10_-transformed JEV RNA copies/ml (milliliter) or g (gram). Colored bars represent subgroups: Control (green bar), litters within Mild clinical subgroup (brown bar), and litters within Severe clinical subgroup (red bar). The dotted horizontal lines represent limit of detection. Viral loads data are also in **S5 Table**. **(K)** JEV negative-strand-RNA-specific RT-PCR data in different tissue samples from piglets. X-axis represents pregnant pig IDs and symbol shapes represent live borne (filled dots) or stillborn (crosses) piglets. Empty dots—fetal membranes (placenta and amnion). Each colored symbol represents different types of tissues: Blue—brains; purple—blood plasma; and black—tonsils. Y-axis represents Ct values. Detailed negative-strand-specific RT-PCR data are in **S6 Table**. Portions of the figure are created in BioRender: Karniychuk, V. (2025) https://BioRender.com/nuv015v and Karniychuk, V. (2026) https://BioRender.com/s8p5mj0.

As expected, a control pig delivered healthy piglets (**Fig 2B**); there were no dead or weak piglets in the litter. Based on clinical outcomes after birth, we subdivided litters from six JEV-inoculated pregnant pigs into two subgroups—mild infection (pigs #166, #167, and #168) and severe infection (pigs #169, #170, and #171) (**Fig 2B**). Three pigs in the mild infection subgroup delivered 48 apparently healthy piglets (89%), only 1 weak (2%), and 5 (9%) dead piglets (with no apparent developmental abnormalities) out of a total of 54 newborns (**Fig 2B**).

Three pigs in the severe infection subgroup delivered 8 apparently healthy piglets (22%), 1 weak piglet (3%), 6 dead piglets with no apparent developmental abnormalities (17%), and 21 dead fetuses/piglets (58%) with severe pathology (decomposition, edema, mummification) out a total of 36 (**Fig 2B-F**). This division into subgroups, based on clinical outcomes, was also consistent with JEV loads in maternal blood (**Fig 2G**). Specifically, the maternal blood plasma samples from the control pregnant pig (pig #164) were negative for JEV; and the three pregnant pigs (pigs #169, #170, and #171) that delivered the severe piglet subgroup (**Fig 2B**) had higher JEV loads in the maternal blood plasma than the three pregnant pigs (pigs #166, #167, and #168) that delivered the mild piglet subgroup (**Fig 2G**).

Furthermore, the division into subgroups, based on clinical outcomes, was also consistent with JEV loads in fetal membranes and piglet tissues (**Fig 2H–2J**). The JEV loads in fetal membranes—placenta and amnion—confirmed transplacental infection (**Fig 2H**). In accordance with the clinical outcomes, all tested placental and amnion samples in three litters with severe clinical outcomes showed high JEV loads (pigs #169–#171; **Fig 2H** and **S5 Table**), while in the mild subgroup amnions from only one litter (out of three) showed positive JEV loads at the limit of detection (pig #167; **Fig 2H** and **S5 Table**). Also, all tested brain samples from dead piglets in severe subgroup had high JEV loads (**Fig 2I** and **S5 Table**), suggesting *in utero* acquisition of JEV that led to subsequent death during fetal period or during birth (**Fig 2C**).

In accordance with the clinical outcomes, all tested nasal swab and tonsil samples from live piglets in litters with severe outcomes showed high JEV loads (**Fig 2J** and **S5 Table)**. In contrast, in the mild subgroup, one live piglet from each litter (pigs #166–#168) had JEV loads in nasal swabs. Also, one live piglet in mild subgroup litter (pig #166) had JEV in tonsils (**Fig 2J** and **S5 Table**). Interestingly, one live piglet from severe subgroup litter (pig #169) had high JEV load (5.3 log_10_ RNA copies/ml) in blood plasma (**Fig 2J** and **S5 Table**), which was comparable to peak JEV RNA loads in most maternal samples (**Fig 2G**).

Next, we tested different tissue samples to confirm productive JEV infection by negative-strand specific RT-PCR, a technique commonly used to confirm active flavivirus replication [25, 26]. JEV negative-strand RNA was detected in fetal membranes—placenta and amnion—in three litters with severe clinical outcomes (pigs #169–#171; **Fig 2K** and **S6 Table**), suggesting productive transplacental infection. Likewise, the brain tissue samples from dead piglets in these three litters (pigs #169–#171) were positive for JEV negative-strand RNA (**Fig 2K** and **S6 Table**), suggesting death *in utero* or during birth because of productive JEV infection. Interestingly, in one liveborn piglet from the severe subgroup litter (pig #169), both tonsil and blood plasma samples showed the presence of JEV negative-strand RNA (**Fig 2K** and **S6 Table**); the Ct value detected in the blood plasma of the piglet (**Fig 2K**; **Table S6**) was comparable to the peak Ct value observed in the corresponding maternal blood (**Table S6**).

We used RNA from the blood plasma samples of JEV-positive mothers (**Fig 2G** and **S5 Table**) for NGS. Library construction and NGS were successful from the blood plasma samples of 4 adult pregnant pigs (pigs #167, #169, #170, and #171 **Fig 3A**) out of 6 pigs from the mild and severe infection subgroup. In these pregnant pigs (pigs #167, #169, #170, and #171 **Fig 3A**), median frequencies of blood plasma mutations, ranged from 5–9% (**Fig 3A**). The percentage of *de novo* emerged iSNVs (triangle symbols in **Fig 3A**) calculated from the total number of sample-specific mutations in blood from pregnant pigs was: pig #167—64%, pig #169—43%, pig #170—0%, and Pig #171—25% (**Fig 3A** and **S7 Table**).

**Fig 3.**
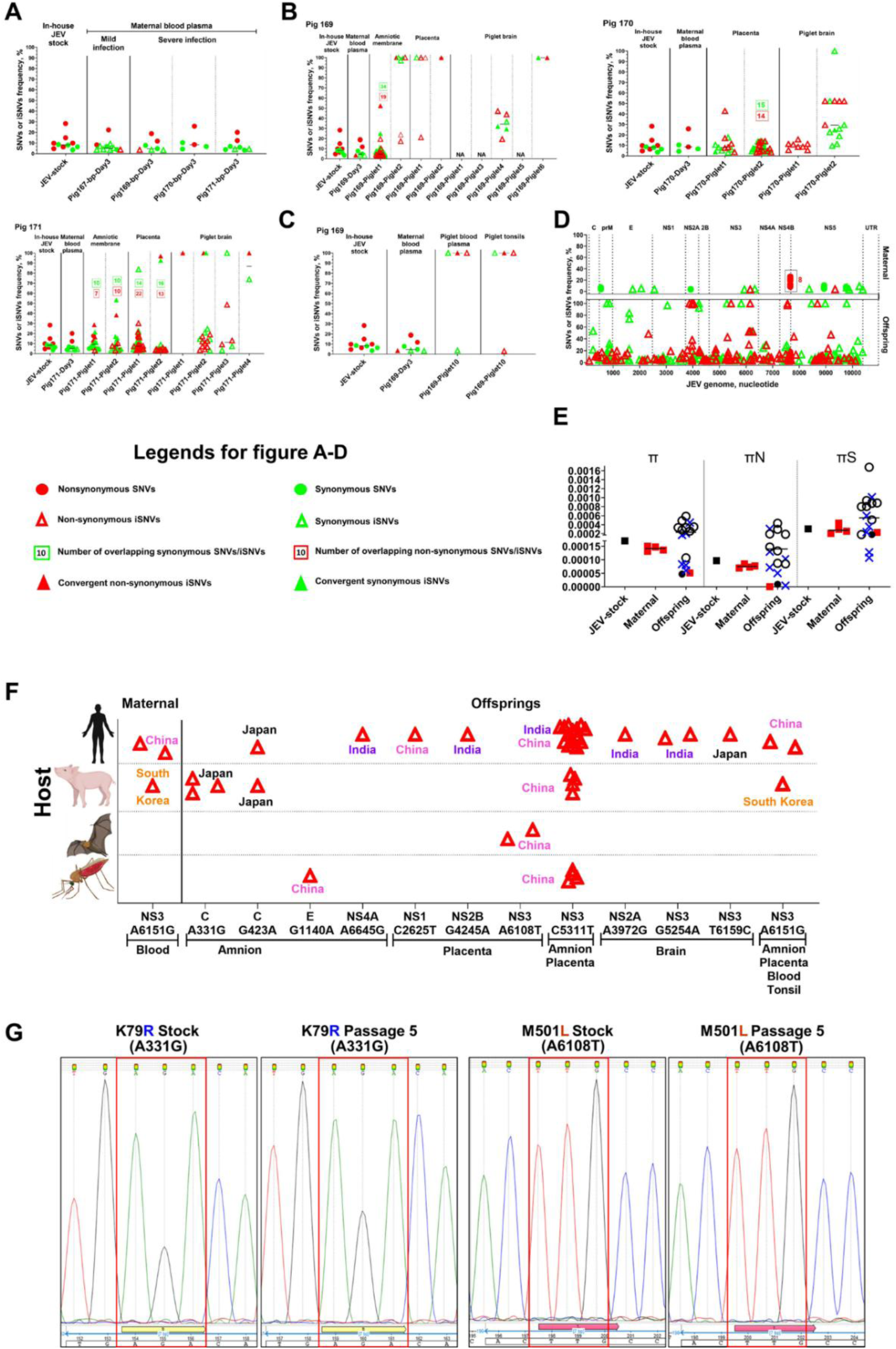
Offspring-specific JEV evolution in the native amplifying host. **(A)** NGS data from maternal-blood plasma collected from pregnant pigs (#167, #169, #170, and #171) on day 3 post JEV inoculation. (**B** and **C**) NGS data from maternal-blood plasma from pregnant pigs (#169, #170, and #171), their fetal membranes expelled after delivery (amnion, placenta), and tissues from piglets (brain, blood plasma, and tonsils). Fetal membrane and piglet tissue samples were collected after birth; the total duration of pregnancy in pigs is 114–116 days. **SNVs:** Single nucleotide variants. **iSNVs:** *de novo* emerged intra-host SNVs. Solid horizontal line represents median values. Details of all SNVs and iSNVs data are in **S7 Table**. (**D**) SNV distribution pattern in the JEV genome. C: JEV genomic region encoding capsid protein; prM: precursor membrane protein; E: envelope protein; NS1: nonstructural protein 1; NS2A: nonstructural protein 2A; NS2B: nonstructural protein 2B; NS3: nonstructural protein 3; NS4A: nonstructural protein 4A; NS4B: nonstructural protein 4B; NS5: nonstructural protein 5. UTR: untranslated region. Raw data are shown in **S7 Table**. (**E**) Nucleotide diversity metrics. X-axis represents sample origin. Black square—JEV stock. Red square—blood plasma. Other symbol shapes represent liveborn (filled dots) or stillborn (crosses) piglets. Empty dots—fetal membranes (placenta and amnion). Each colored symbol represents different types of tissues: Blue—brains; and black—tonsils. Y-axis: Mean pairwise difference for π (mean number of pairwise differences per site across the whole genome), πN (mean number of pairwise non-synonymous differences per non-synonymous site across the genome) and πS (mean number of pairwise synonymous differences per synonymous site across the genome). Solid horizontal line represents median values. Raw data are shown in **S7 Table**. (**F**) Comparative experimental (maternal and offspring; **S7** and **S8 Tables**) and field JEV evolution. Maternal and offspring *de novo* emerged non-synonymous iSNVs identified in this study and previously reported in field studies (in the NCBI database; see supplementary materials and methods and **S8 Table**). Red empty triangles represent previously reported field occurrences of iSNVs identified in this study. X-axis represents specific iSNV and their position within the JEV genome. C: JEV capsid protein. pr-M: Precursor membrane protein. E: Envelope protein. NS1: Nonstructural protein 1. NS2A: Nonstructural protein 2A. NS2B: Nonstructural protein 2B. NS3: Nonstructural protein 3. NS4A: Nonstructural protein 4A. NS4B: Nonstructural protein 4B. NS5: nonstructural protein 5. UTR: untranslated region. (**G**) Sanger sequencing. Offspring *de novo* emerged non-synonymous iSNVs A331G (amino acid substitution K79R) and A6108T (amino acid substitution M501L) identified in this study and previously reported in the field studies (**S7** and **S8 Tables**). Introduced substitutions showed stability after 5 passages in Vero cells. Nucleotide peaks highlighted in red squares encode introduced K79R and M501L amino acid substitutions. Raw FASTQ NGS files are deposited in BioProject: #PRJNA1467537. Portions of the figure are created in BioRender: Karniychuk, V. (2026) https://BioRender.com/xw9t9tk.

For piglet samples from all 3 litters (from pigs #169, #170, and #171), the percentage of piglet SNVs in different sample (amnion, placenta, brain, tonsils, blood plasma) (solid colored dots in **Fig 3B** and **3C**) calculated from the total number of sample-specific mutations was considerably lower than the percentage of iSNVs (solid colored triangles in **Fig 3B** and **3C**): amnion—SNVs 7% versus iSNVs 93%, placenta—2% versus 98%, brain—0% versus 100%, tonsils—0% versus 100%, and blood plasma—0% versus 100%.

For pig #167, #169, #170, and #171, maternal and offsprings iSNVs were located in the JEV genomic regions encoding all structural (prM, C, and E) and non-structural (NS1, NS2A, NS2B, NS3, NS4A, NS4B and NS5) proteins (**Fig 3D**).

To quantify the degree of convergent evolution in the JEV genome, any iSNV that emerged in more than one piglet was defined as convergent, and the total convergence was quantified as the percentage of convergent iSNVs relative to the total number of iSNVs. In litters from pig #169, the degree of convergent evolution was 9%, synonymous iSNVs—4%, and non-synonymous iSNVs—5% (**Fig 3D** and **S7 Table**). In litters from pig #170, the degree of convergent evolution was 3%, synonymous iSNVs—1.5%, and non-synonymous iSNVs—1.6% (**Fig 3D** and **S7 Table**). In litters from pig #171, the degree of convergent evolution was 9%, synonymous iSNVs—6%, and non-synonymous iSNVs—3% (**Fig 3D** and **S7 Table**). Also, we found a convergent non-synonymous iSNVs (C5311T corresponding to A235V amino acid substitution) from the litters from two different pigs (pigs #169 and #170). Interestingly, a convergent non-synonymous iSNV (A6151G corresponding to E515G amino acid substitution) was observed within NS3 region of the JEV genome in both maternal-blood plasma and tissues (amnion, placenta, blood plasma, and tonsils) of piglets from pig #169 (**Fig 3B, 3C** and **S7 Table**). The frequency of this specific non-synonymous iSNV was 3.6% in maternal blood. However, it showed a considerable increase, rising to 100% in the piglet tissues (amnion, placenta, blood, and tonsils). This increase may suggest distinct maternal and fetal evolutionary forces and an evolutionary placental bottleneck during transplacental transmission, as the other 6 maternal-blood plasma non-synonymous SNVs and iSNVs (**Fig 3B** and **3C**) were not present in the samples collected from piglets after birth.

Like in the fetal study (**Fig 1F**), the low nucleotide diversity (π) across maternal and offspring samples (**Fig 3E**) suggests that overall viral populations remain genetically constrained within both biological compartments. πS also exceeded πN in both compartments (**Fig 3E**), consistent with purifying selection [24]. However, despite the overall low diversity, the consistent emergence of compartment-specific iSNVs in amniotic membranes, placenta, brain, and tonsils (**Fig 3A-E**) indicates ongoing localized virus evolution and the emergence or enrichment of potentially biologically meaningful JEV variants.

We compared iSNVs that emerged during JEV infection in pregnant pigs and offspring (**S7 Table**) with JEV sequences available in the NCBI nucleotide database. For maternal samples, one *de novo* emerged non-synonymous iSNVs (A6151G corresponding to E515G amino acid substitution) out of 12 total identified iSNVs was reported during 1949-2017 in JEV sequences from China and South Korea and identified in field samples from humans and pig (**Fig 3F** and **S8 Table**).

In offspring samples, 12 *de novo* emerged non-synonymous iSNV (**Fig 3F** and **S8 Table**) out of 263 total identified iSNVs were reported previously during 1935-2019 in JEV sequences from Japan, China, India, and South Korea. These offspring iSNVs were previously identified in field samples from human blood and cerebrospinal fluid, mosquitoes, pig sera, and bats (**Fig 3F** and **S8 Table**). We used reverse genetics to synthetically rescue JEV variants containing the A331G (K79R amino acid substitution) and A6108T (M501L amino acid substitution) mutations (the reason for selecting these two mutations is provided in **S1 File**). Both variants were passaged five times in cell culture, and stability of the mutations was confirmed by Sanger sequencing (**Fig 3G**); the parental JEV variant retained the wild-type sequence before and after passaging.

Altogether, the JEV genetic heterogeneity acquired during fetal infection (**Fig 1C-E**) persists in expelled fetal membranes (amniotic and placental membranes) and demised (piglet brain) or surviving (piglet blood plasma and tonsils) offspring (**Fig 3A-D**).

### Exposure to JEV during fetal life causes transcriptional footprints in offspring blood cells and tonsils

Global host mRNA expression in samples from the offspring study (**Fig 2A**) was evaluated by RNA-seq. We used whole blood cells and tonsil tissues from 9 piglets from the Control subgroup, 24-36 (whole blood cells, n=24; tonsils, n=36) piglets from the Mild clinical subgroup, and 9 survived piglets from the Severe clinical subgroup (**Fig 2B** and **S9 Table**).

First, we compared global gene expression (**Fig 4**) in offspring whole blood cells from Control versus Mild clinical subgroups. JEV-affected offspring had profound gene transcriptional changes: 613 downregulated and 215 upregulated genes (**Fig 4A** and **S10 Table**). Significantly affected, differentially expressed genes (DEGs) in whole blood cells were further analyzed by comparing these DEGs (**Fig 4A**) to a set of 48 pig or 271 human genes known to be involved in the interferon (IFN) signaling pathway (**S11 Table**). For the pig IFN gene set, the analysis showed 7 downregulated and 1 upregulated gene; for the human set 36 downregulated and 3 upregulated genes (**Fig 4A; S10 Table**). All IFN genes identified in the pig set were present in the human gene set.

**Fig 4.**
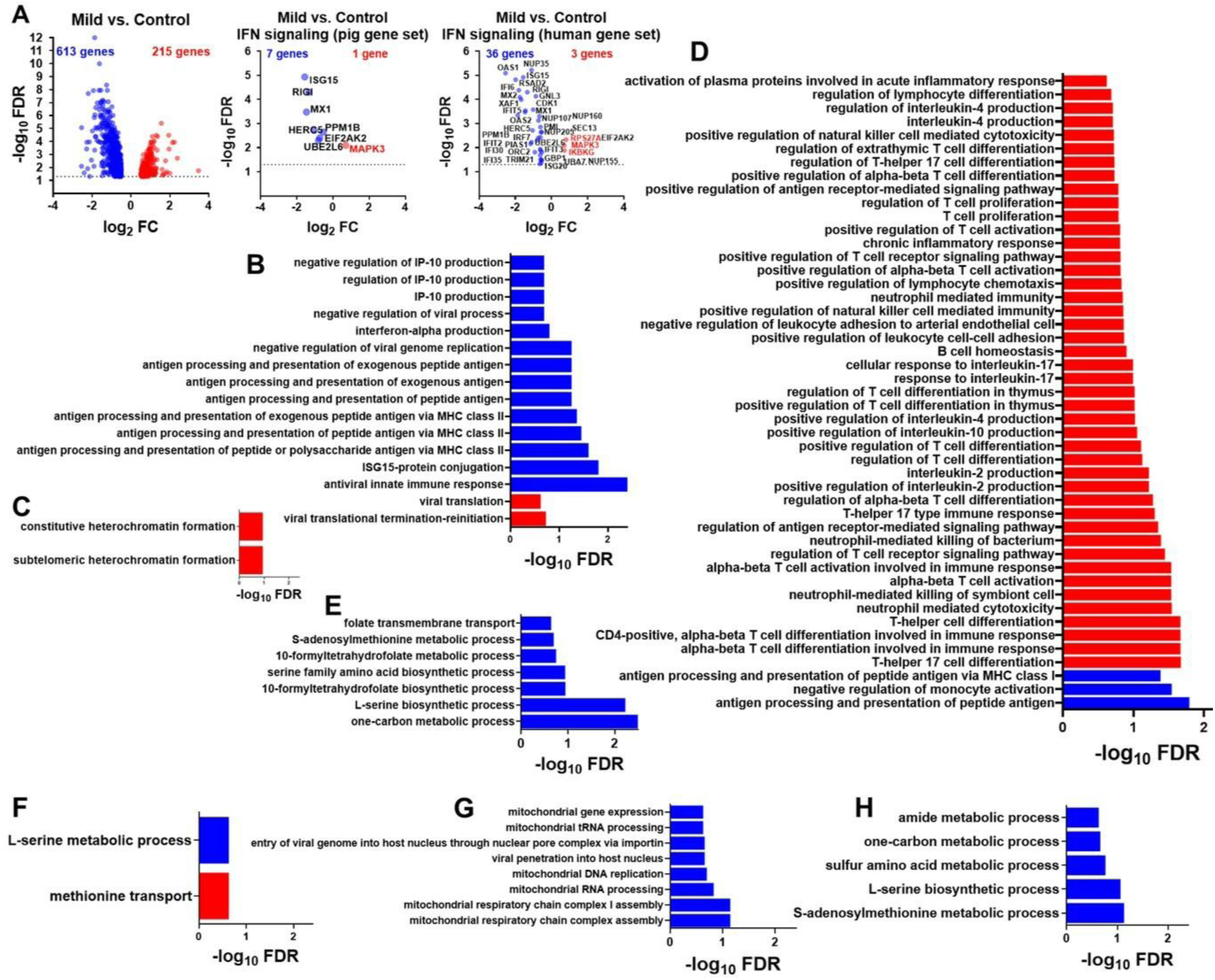
Transcriptional pathology in whole blood cells and tonsils of offspring affected by JEV during fetal life. (**A**) Upregulated (red) and downregulated (blue) genes in whole blood cells from the Mild clinical subgroup (raw data in **S10 Table**). Upregulated and downregulated genes related to IFN signaling (pig or human gene sets; **S11 Table**) in the Mild clinical subgroup are also shown (**S10 Table**). Direction of DEG analysis: Mild clinical subgroup/Control group. Genes with FDR < 0.05 and log_2_ fold change (FC) ≥ 0.5 (1.41-fold) are shown. FDR: false discovery rate. Significantly altered (FDR-adjusted *P* < 0.25) GO biological processes in the Mild clinical subgroup whole blood cells related to viral infection and immune responses (**B**) and epigenetic memory (**C; S10 Table**). Significantly altered GO biological processes in the Mild clinical subgroup tonsils related to T cell responses (**D**) and epigenetic memory (**E; S16 Table**). Significantly altered GO biological processes in the Severe clinical subgroup whole blood cells (**F**) and tonsils (**G**, **H**) related to mitochondrial function/dysfunction (**G**) and epigenetic memory (**F**, **H; S17** and **S18 Tables**). Raw FASTQ NGS files are deposited in BioProject: #PRJNA1467734.

Along with profound gene transcriptional changes, JEV-affected offspring had 124 significantly affected GO biological processes (**S10 Table**). After narrowing the analysis to GO processes related to viral infection and immune responses, genes with significantly upregulated expression in whole blood cells were enriched for GO processes related to “viral translational termination-reinitiation” and “viral translation” (**Fig 4B**). Also, in accordance with downregulated DEGs related to IFN signaling (**Fig 4A**), 12 GO processes related to IP-10 production, IFN-alpha regulation, antigen presentation, and antiviral innate immune response were negatively enriched (**Fig 4B**). The lowest *P*-value (FDR = 0.0003) in whole blood cells was observed for the downregulated GO biological process “antiviral innate immune response.”

Second, because the number of piglets and whole blood cell samples in the Mild clinical subgroup (24 whole blood cell samples) was substantially higher than in the Control group (9 samples) or the Severe clinical subgroup (9 samples), we compared global gene expression in offspring whole blood cells from the Control group with whole blood cells from 7–9 offspring in each Mild subgroup litter (litters: #166, #167, and #168; **S9 Table**; **S12-14 Tables**). The comparative analysis confirmed that whole blood cells from each litter had 3–8 (pig IFN gene set) and 17–34 (human set) affected DEGs related to IFN signaling (**S15 Table**). Many IFN signaling genes and GO processes overlapped between three outbred litters (**S10 Table** and **S12-15 Tables**), including negatively enriched “antiviral innate immune response” GO process, showing that exposure to JEV during fetal life causes transcriptional footprints in offspring blood cells in each outbred litter within the Mild subgroup.

Interestingly, among affected GO processes in whole blood cells were “subtelomeric heterochromatin formation” and “constitutive heterochromatin formation” (**Fig 4C**; **S10 Table**) which are involved in a core mechanism of epigenetic memory.

Next, we compared global gene expression in offspring tonsils from Control versus Mild clinical subgroups. JEV-affected offspring had 20 downregulated and 24 upregulated genes (**S16 Table**). These DEGs in tonsils were further analyzed by comparing to a set of 48 pig or 271 human genes involved in the IFN signaling pathway (**S11 Table**). The analysis showed 3 significantly downregulated genes related to IFN signaling— XAF1, OAS2, and OAS1 (**S16 Table**). To further extend analysis beyond DEGs, we conducted Gene Set Enrichment Analysis (GSEA) analysis which offers high sensitivity due to statistical rigor where each gene in the entire input set (**S16 Table**), not only genes with FDR < 0.05 and log2 fold change (FC) > 0.5, is statistically processed and used for annotation [27, 28]. JEV-affected offspring had 672 affected GO biological processes in tonsils (**S16 Table**). Relevant to the lymphoid organ function of tonsils and their critical role in the T cell response, many GO processes related to T cells and T cell interleukins were positively enriched (**Fig 4D**). Processes related to other immune cell types—neutrophils and B cells—were also affected. Interestingly, among positively enriched GO processes were “chronic inflammatory response” and “activation of plasma proteins involved in acute inflammatory response” (**Fig 4D**). Among the negatively enriched GO processes in tonsil samples were 7 processes that have a strong direct influence on epigenetic memory (**Fig 4E**), specifically by regulating S-adenosyl-L-methionine (SAM) production, the universal methyl donor for DNA methyltransferases and histone methyltransferases.

Next, we compared global gene expression in offspring whole blood cells and tonsils from Control versus Severe clinical subgroups. JEV-affected offspring from the Severe subgroup had 12 downregulated and 11 upregulated genes in whole blood cells (**S17 Table**). Only one DEG was related to IFN signaling—OAS1 (**S17 Table**). More sensitive GSEA analysis however revealed “viral translation” (positively enriched) and “antiviral innate immune response” (negatively enriched) significantly affected processes (**S17 Table**).

While DEGs were not identified in tonsils after comparing global gene expression in Control versus Severe clinical subgroups, more sensitive GSEA analysis, which statistically processes the entire input set with more than 19 thousand transcripts [27, 28] (**S18 Table**), showed 121 affected GO biological processes (**S18 Table**). Among negatively enriched GO processes were “viral penetration into host nucleus” and “entry of viral genome into host nucleus through nuclear pore complex via importin” (**Fig 4G**), which is consistent with persistent JEV infection in tonsils (**Fig 2J** and **2K**) and known flavivirus interactions with the importin and host cell nuclear [29, 30]. Interestingly, six GO processes related to mitochondrial function were negatively enriched (**Fig 4G**) suggesting a lasting impact on cellular energy production, potential mitochondrial dysfunction including long-lasting epigenetic changes [31–34]. This may reflect the persistent JEV infection in tonsils (**Fig 2J** and **2K**), as well as the presence of long living tissue-resident memory T cells that may sustain acquired long-lasting transcriptional pathology in tonsils.

Among affected GO processes in whole blood cell (**Fig 4F** and **S17 Table**) and tonsil (**Fig 4H** and **S18 Table**) samples from the Severe clinical subgroup, were several processes which are known to have strong direct influence on epigenetic memory via SAM production, including the common GO process—“L-serine metabolic process.”

The interesting common finding is that altered transcriptional responses in all available samples—whole blood cells and tonsils, from all clinical subgroups (Mild and Severe)—included affected epigenetic processes and memory (described above). We previously found that transplacental JEV infection in pregnant pigs is associated with increased IFN*α* levels in live JEV-exposed fetuses [12]. All control fetuses had no detectable IFN*α* or levels at the limit of detection in their blood [12]. Our working hypothesis is that during JEV infection in pregnant pigs, surviving fetuses that were exposed *in utero* to inflammatory cytokines, including IFN*α*, may carry long-lasting immunopathology at least partially determined by epigenetic alterations. The hypothesis is indirectly supported by RNA-seq data in JEV-affected offspring (**Fig 4**), and by functional cellular sequelae that we previously identified in offspring affected by Zika virus, the flavivirus related to JEV [35]. Moreover, the general harmful effects of excessive IFN*α* exposure and its contribution to immune exhaustion, immunosuppression, and immunopathology are well documented (not in fetuses and offspring) [36, 37]. To further test the hypothesis, we employed a reductionist model in which fetal CD34+ hematopoietic stem progenitor cells (HSPCs) obtained from umbilical cord blood were exposed to IFN*α* during differentiation to identify transcriptional and functional footprints in HSPC-derived granulocyte-macrophage progenitors. Also, to define whether IFN*α*-induced molecular footprints can persist in CD34+ HSPC-derived differentiated granulocyte-macrophage progenitors, we used whole-genome methylation to profile epigenetic modifications. The experimental setup is represented in **S3 Fig**.

First, to partially reproduce a fetal environment with excessive IFNα, we treated fetal CD34+ HSPCs derived from the same donor with recombinant IFNα or Control media for 24 h. Afterward, for the differentiation assay, IFNα-treated (for 24 h) CD34+ HSPCs were cultured in a complete methylcellulose medium for 10 days in the presence or absence of IFN-α that resulted in two conditions—IFNα-short-treated (24 h before differentiation) and IFNα-long-treated cells (24 h before differentiation + 10 days during differentiation) (**S3 Fig)**. For the Control condition, CD34+ HSPCs were not exposed to IFNα at any experimental stage. After 10 days, we quantified granulocyte-macrophage progenitors differentiated from CD34+ HSPCs in both IFNα and Control conditions and compared their whole-genome transcription by RNA-seq (**Fig 5**).

**Fig 5.**
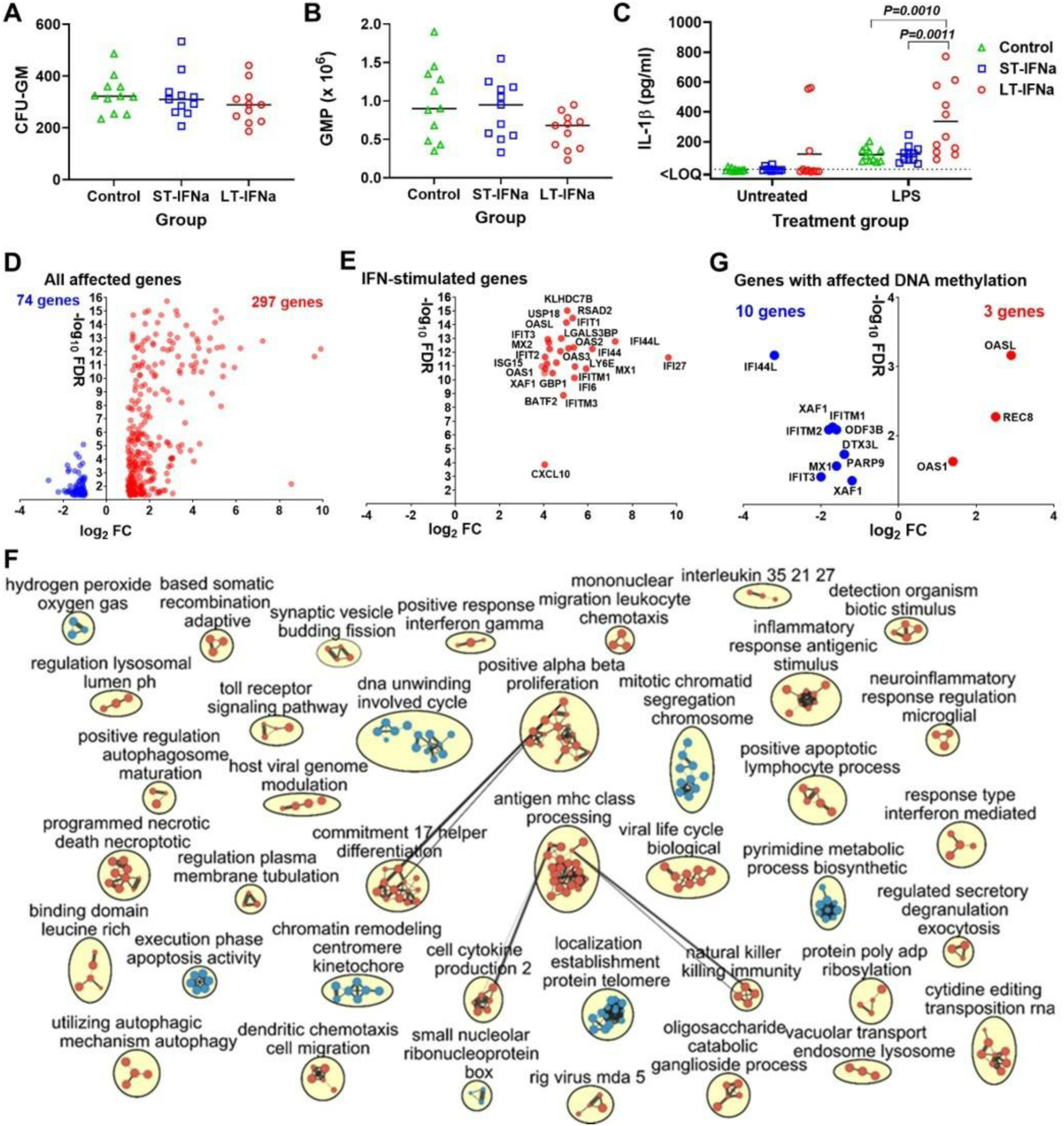
Transcriptional, epigenetic, and functional alterations in fetal hematopoietic stem cells differentiating during IFNα exposure. (**A** and **B**) Proliferation and differentiation of fetal CD34^+^ HSCs during IFNα exposure. The absolute number of granulocyte-macrophage colony-forming units (**CFU-GM**) (**A**) and granulocyte-macrophage progenitors (**GMP**) (**B**). ST-IFNα: IFNα-short-treated. LT-IFNα: IFNα-long-treated. (**C**) Lipopolysaccharide (LPS)-induced IL-1*β* response in GMP differentiated from CD34+ HSCs in different IFNα treatment condition. Solid lines represent the mean. The dotted line represents the limit of quantification (LOQ). The difference with *P* < 0.05 was considered statistically significant. IL-1*β* levels were compared between different cellular conditions with Kruskal–Wallis H-test. (**D** and **E**) Transcriptional (RNA-seq) responses in GMP differentiated from CD34+ HSPCs in the long-term IFNα treatment condition. Upregulated (red) and downregulated (blue) genes. All affected genes (**D**) and the top 25 genes encoding canonical IFN-stimulated genes **(E)** with FDR-adjusted *P* < 0.05 and log_2_ fold change (FC) > 1 are shown. Raw data are in **S19 Table**. (**F**) An enrichment map of significantly altered GO biological processes in GMPs differentiated from CD34+ HSPCs in the long-term IFNα treatment condition under RNA-seq analysis. Red are pathways with positive and blue are with negative enrichment. All subnetworks with FDR-adjusted *P* < 0.25 and at least three connected nodes are shown. Raw data are in **S19 Table**. Direction of analysis: IFNα-exposed cells / Control cells. An accession number for RNA-seq data is PRJNA836933 in NCBI BioProject. (**G**) Genes with different DNA methylation and mRNA expression identified in GMPs differentiated from fetal CD34+ HSPCs in the long-term IFNα treatment condition. Raw data are in **S20 Table**. Direction of analysis: IFNα-exposed cells / Control cells. Raw data from the methylation microarray with the Infinium MethylationEPIC BeadChip Kit (Illumina) and Illumina iScan are in DRYAD: https://doi.org/10.5061/dryad.v6wwpzhbq

The colony-forming unit assay—the commonly used assay to measure the proliferation and differentiation of HSPCs in humans and mice [38]—showed that fetal CD34+ HSPCs exposed to IFNα for the short-term have the same number of granulocyte-macrophage colony-forming units and individual granulocyte-macrophage progenitors as other experimental conditions (**Fig 5A** and **B**). However, CD34+ HSPCs exposed to IFNα for the long term had a trend to the lower number of individual granulocyte-macrophage progenitor cells (**Fig 5B**). While the difference was not statistically significant (*P* > 0.05), the trend suggests altered proliferation and differentiation of HSPCs in IFN-α-long-treated condition.

In progenitors differentiated in the short-term IFNα treatment condition, individual gene analysis showed no significant transcriptional changes (**S19 Table**). However, more sensitive GO biological-process enrichment analysis by GSEA identified 26 processes related to mitochondrial function/dysfunction, type I IFN, and regulation of the viral cycle (**S19 Table**). Interestingly, among affected GO processes was “positive regulation of histone H3-K4 methylation,” a critical epigenetic process associated with active chromatin and transcriptional memory.

Progenitors differentiated in the long-term IFNα treatment condition had profound transcriptional changes with 297 upregulated and 74 downregulated genes (FDR-adjusted *P* < 0.05, log_2_ FC > 1; **Fig 5D** and **S19 Table**). Among the top thirty upregulated genes with 4-10 log_2_ fold change were 25 genes encoding canonical ISGs (**Fig 5E** and **S19 Table**). Accordingly, enrichment of GO biological processes showed significant effects in granulocyte-macrophage progenitors treated with IFNα for a long time with 317 upregulated and 132 downregulated processes (FDR-adjusted *P* < 0.25; **Fig 5F** and **S19 Table**). Genes with altered expression were positively enriched for processes related to interferon responses (15 GO processes FDR-adjusted *P* < 0.25), inflammatory responses (12 GO processes), virus-host interactions (23 GO processes), cellular immune responses, antigen processing (88 GO processes), and cell death (23 GO processes) (**Fig 5F** and **S19 Table**). Several affected processes, including “ATP dependent chromatin remodeling,” “histone H2B ubiquitination,” “DNA replication independent nucleosome organization,” (**S19 Table**) were directly linked to epigenetic memory via regulating chromatin accessibility, stability, and nucleosome positioning.

Next, to define potential functional alterations, we quantified and compared IL-1*β* responses after LPS stimulation in granulocyte-macrophage progenitors differentiated from CD34+ HSCs in the short-term IFNα treatment, long-term IFNα treatment, and Control conditions. Granulocyte-macrophage progenitors differentiated in the Control and short-term IFNα conditions showed on average 7 to 10-fold increases in IL-1*β* concentrations after LPS stimulation compared to the background concentrations in not stimulated with LPS progenitors (**Fig 5C**). Progenitors from 3 donors in the long-term IFNα treatment condition showed increased IL-1*β* concentrations even with no LPS stimulation (**Fig 5C**). But when three outliers were removed, LPS stimulation induced an IL-1*β* response with an average 88-fold increase compared to the background concentrations in LPS-not stimulated progenitors (**Fig 5C**). Also, after LPS stimulation, IL-1*β* concentrations in long-term IFNα treated progenitors were significantly higher than in progenitors of the Control (*P* = 0.0010) and short-term (*P* = 0.0011) IFNα conditions. Transcriptional data also confirmed affected genes related to the Toll-like receptor 2 signaling pathway (**S19 Table**) that may alter LPS responses [39, 40]. Inflammasomes produce IL-1*β*; accordingly, genes of main inflammasome components [41], including sensors (NLRP1, NLRP3, IFI16) and effector enzymes (CASP1, CASP4, CASP5) were significantly affected (**S19 Table**).

Finally, we analyzed genome-wide DNA methylation in granulocyte-macrophage progenitors differentiated from CD34+ HSPCs in the Control and long-term IFNα treatment conditions. 33 genes were differentially methylated in the long-term IFNα treatment condition (FDR-adjusted *P* < 0.05; log_2_ FC > 1; **S20 Table**). Among these differentially methylated genes, 13 genes were also differentially expressed as identified by RNA-seq (**Fig 5G** and **S20 Table**). Ten commonly affected genes were ISGs. Ten genes had decreased DNA methylation and increased mRNA expression (**Fig 5G**). Enrichment of GO biological processes based on different DNA demethylation showed significant effects in granulocyte-macrophage progenitors treated with IFNα, with 31 affected processes (FDR-adjusted *P* < 0.05) related to type I IFN responses, defense responses, innate immunity, and viral life cycle (**S20 Table**). Eleven GO processes were affected in both whole-genome methylation and gene expression RNA-seq analysis: “response to interferon-alpha,” “response to interferon-beta,” “response to type I interferon,” “response to interferon-gamma,” “defense response to virus,” “response to virus,” “regulation of viral genome replication,” “negative regulation of viral genome replication,” “viral life cycle,” “negative regulation of viral life cycle,” and “negative regulation of viral process” (**S20 Table**).

Altogether, exposure to JEV during fetal life induces transcriptional footprints in blood cells and tonsils of live offspring from both Mild and Severe clinical subgroups, including alterations in pathways associated with epigenetic regulation and memory. In addition, we found that both short-term and, particularly, prolonged exposure to IFNα affects the differentiation of fetal CD34+ HSPCs, resulting in transcriptional, functional, and epigenetic changes in HSPC-derived granulocyte–macrophage progenitors. Collectively, these *in vitro* and *in vivo* findings provide directions for future studies investigating the mechanisms underlying molecular footprints and long-term sequelae in JEV-affected offspring.

## Discussion

The identified prolonged persistence of JEV in fetuses, which continues into the postnatal period in offspring, may contribute to maintenance of the JEV transmission cycle. Non-vector transmission of JEV has been defined both in the field and under experimental conditions and represents veterinary and public health concerns [9, 19, 42]. Mathematical modeling based on longitudinal data from pigs on JEV-endemic farms in Cambodia supports direct non-vector transmission between pigs, which may be particularly important for transmission during seasons with few or no mosquitoes [43]. Based on our findings, we hypothesize that JEV may exploit prolonged months-long persistence in fetuses as a natural mechanism to bridge inter-mosquito seasons on swine farms and in wild pig populations. Indeed, the duration of pig gestation (∼114 days), together with the experimentally defined window of susceptibility to transplacental and fetal JEV infection (days 30-80 of pregnancy in our previous study [12] and from 50 days to birth at 114-116 days in the present study), could at least partially bridge the 2-4-month period of reduced mosquito activity observed in for example Cambodia or other JEV-endemic regions.

There are several potential pathways via which fetal persistent JEV may be shed after birth. First, the expelled placenta, fetal membranes, and dead fetuses represent a likely source. We detected high viral RNA loads in afterbirth placentas, amnion, and fetal brains (**Fig 2H**), including negative-strand JEV RNA (**Fig 2K**), suggesting active replication. Placental tissues may be consumed (placentophagy) by pigs on farms and particularly in wilderness by pigs, predators, or opportunistic scavengers. Moreover, under typical farm practices, placenta, fetal membranes, and stillborn piglets are often disposed of in outdoor compost piles consisting of carcasses and layers of carbon material. This creates a potential route for environmental contamination, including mosquito breeding sites. Notably, flavivirus transmission to mosquitoes from contaminated breeding sites including sewage has been demonstrated [44]. Second, one piglet in this study had high JEV RNA loads in blood (**Fig 2J**) including negative-strand JEV RNA (**Fig 2K**), representing a measurable fraction of the sampled population (1.7%; one blood JEV-positive piglet out of 58 liveborn piglets). Considering the scale of swine production in Asia and the presence of potentially similar dynamics in wild boar populations, even low-frequency events may represent a meaningful source of transmission, enabling *in utero*-acquired JEV to infect mosquitoes and subsequently spread to other hosts. Finally, tonsils appear to represent a site of persistent JEV infection in affected offspring (**Figs 2J** and **2K**). JEV persistence in tonsils has previously been demonstrated in experimentally infected young piglets and adult pigs [9, 12, 18, 19] and has been associated with weak induction of innate and adaptive immune responses [45]. Viruses persisting in tonsils can be shed in saliva and nasal fluids [9, 12, 18, 19, 46, 47], which, together with JEV detected in nasal swabs (**Fig 2J**), may contribute to postnatal JEV transmission to naïve piglets via direct contact. Interestingly, JEV RNA identified in offspring tonsils and nasal swabs in the present study (**Fig 2J**) were comparable to those reported in tonsils and oronasal swabs in the pioneering study that established direct-contact JEV transmission between experimentally infected young piglets [19].

Genomic heterogeneity across maternal, fetal, and offspring tissues (**Figs 1** and **3**), suggests complex viral diversity within interconnected yet distinct biological compartments. Recently, the strong positive selection of two JEV mutations was reported during experimental passage in young piglets and porcine macrophages [48, 49]. Interestingly, fetal infection may represent an additional source of JEV genetic diversification and variant emergence, as indirectly supported by comparisons with publicly available JEV sequences (**Figs 1** and **3**). This observation points to a new conceptual aspect of JEV evolutionary biology, distinct from the traditionally described evolutionary mechanisms in young and adult mammals, avian hosts, and mosquito vectors.

Here, for the first time, we demonstrate transcriptional postnatal sequelae caused by JEV infection in the natural amplifying host, revealing alterations in clinically normal (visually healthy) piglets. The extent of transcriptional pathology, reflected by a higher number of DEGs and affected GO pathways, was greater in the Mild clinical subgroup than in the Severe clinical subgroup (**Fig 4**). The working theory is that the Mild clinical subgroup reflects a more recent and/or relatively localized transplacental JEV infection, as suggested by limited viral detection in only several amniotic tissues (**Fig 2H**), nasal swab samples and tonsils (**Fig 2J**). Such later and/or localized persistent infection may trigger a cascade of signaling events broadly affecting gene expression in local virus-positive and distant virus-negative tissues. A similar gene-expression pattern was reported in a recent JEV study, where early infection restricted to a very small region of the mouse brain induced widespread transcriptional changes across the entire brain including JEV-negative brain regions [50]. In contrast, the Severe clinical subgroup likely reflects earlier fetal infection with extensive viral dissemination in fetuses, leading to the death of all (**Fig 2B**; litters #170 and #171) or 47% (**Fig 2B**; litter #169) of fetuses, strong activation of immune gene expression followed by JEV-mediated suppression of numerous genes in survived fetuses, which our sampling window did not capture, and ultimately exhaustion (**Fig 4**) of transcriptional responses in newborn piglets. While hypothetical, immune activation in fetal HSPCs exposed to IFNα (transplacental JEV infection in pregnant pigs increases IFNα levels in live exposed fetuses [12]) may partially model activation of immune gene expression in fetuses that our *in vivo* sampling window did not capture. The observed transcriptional postnatal sequelae extend the significance of fetal JEV infection beyond developmental pathology, economic losses to the swine industry, and potential contribution to JEV transmission. The birth of visually healthy but JEV-affected piglets that remain in production systems may increase population-level susceptibility to secondary infections; for example, to zoonotic influenza A virus, Nipah virus, and *Streptococcus suis*.

Collectively, we demonstrate that JEV infection in fetuses of an amplifying porcine host persists postnatally and induces lasting transcriptional footprints in offspring. Our findings provide a strong experimental framework for several novel hypotheses in JEV biology, including: (i) potential transmission pathways linking fetal infection to offspring and subsequent environmental dissemination; (ii) fetal infection as an additional source of JEV genetic diversification and variant generation; and (iii) fetal exposure to JEV as a determinant of increased offspring susceptibility to zoonotic pathogens and potentially enhanced spillover efficiency.

## Acknowledgments

We thank Vaccine and Infectious Disease Organization animal care technicians and veterinarians for their help with animal experiments.

## Data Availability

All relevant data are within the manuscript and its supplementary files. Accession numbers for NGS and RNA-seq data are PRJNA1467537, PRJNA1467734, PRJNA836933, in NCBI BioProject. Raw data from the methylation microarray with the Infinium MethylationEPIC BeadChip Kit (Illumina) and Illumina iScan are in DRYAD: https://doi.org/10.5061/dryad.v6wwpzhbq.

## Funding Statement

This work was supported by US Department of Defense, FY20 PRMRP-Discovery Award W81XWH-21-1-0014 and Canadian Institutes of Health Research (CIHR) Project Grant 424307 to UK. PPS received a scholarship from the School of Public Health, University of Saskatchewan. Vaccine and Infectious Disease Organization receives operational funding from the Government of Saskatchewan through Innovation Saskatchewan and the Ministry of Agriculture, and from the Canada Foundation for Innovation via the Major Science Initiatives for its CL3 facility. The funders had no role in the study design, data collection and analysis, or decision to publish.

## Conflicts of Interest

The authors declare that the research was conducted in the absence of any commercial or financial relationships that could be construed as a potential conflict of interest.

## Supplementary data

**S1 Fig. Maternal SNVs and iSNVs (Fetal study).**

Single nucleotide variants in samples from adult pregnant and non-pregnant pigs. SNVs: Single nucleotide variants. iSNVs: *de novo* emerged intra-host SNVs. NA: Library construction for NGS in these samples was not successful. Detailed SNV data are in S1 Table. Solid horizontal lines represent median values.

**S2 Fig. Fetal SNVs and iSNVs (Fetal study).**

Nucleotide position of SNVs and iSNVs within the JEV genome. Pattern of JEV SNVs and iSNVs in the maternal and fetal blood from pregnant Pig A and Pig C, and their fetuses. C: JEV capsid protein. pr-M: Precursor membrane protein. E: Envelope protein. NS1: Nonstructural protein 1. NS2A: Nonstructural protein 2A. NS2B: Nonstructural protein 2B. NS3: Nonstructural protein 3. NS4B: Nonstructural protein 4B. NS5: nonstructural protein 5. UTR: untranslated region. Raw data are shown in S1 Table.

**S3 Fig. The HSPC experimental setup.**

The figure represents testing of CD34+ HSPCs from one donor and technical replicates. Blood from 11 donors was tested in the same manner. HSPCs: Hematopoietic progenitor stem cells. CFU-GM: Granulocyte-macrophage colony-forming units. LPS: lipopolysaccharide.

**S1 File.**

**Supplementary Materials and Methods.**

**S2 File.**

**In-house whole-genome JEV Nakayama reference sequence.**

**S1 Table. SNVs and iSNVs JEV stock, Maternal, and Fetal (Fetal study).**

**S2 Table. JEV viral loads in Maternal and Fetal samples (Fetal study).**

**S3 Table. NGS read depth and coverage (Fetal & Offspring).**

**S4 Table. Comparative Maternal, Fetal, and Field JEV evolution (Fetal study).**

**S5 Table. JEV viral loads in Maternal and Piglet samples (Offspring study).**

**S6 Table. Negative strand qPCR, Maternal and Piglet tissues (Offspring study).**

**S7 Table. SNVs and iSNVs Maternal and Piglets (Offspring study).**

**S8 Table. Comparative Maternal, Piglet, and Field JEV evolution (Offspring study).**

**S9 Table. Sample IDs for RNA-seq (piglets, Whole blood cells and Tonsils) (Offspring study).**

**S10 Table. RNA-seq, Mild clinical subgroup, Combined 3 litters, DEGs, GOs, Whole blood cells (Offspring study).**

**S11 Table. RNA-seq, Pig and Human reference IFN signaling gene sets (Offspring study).**

**S12 Table. RNA-seq, Mild_166, DEGs, GOs, Whole blood cells (Offspring study).**

**S13 Table. RNA-seq, Mild_167, DEGs, GOs, Whole blood cells (Offspring study).**

**S14 Table. RNA-seq, Mild_168, DEGs, GOs, Whole blood cells (Offspring study).**

**S15 Table. RNA-seq, Mild_Common, DEGs between individual 166, 167, or 168 litters and combined 3 litters, Whole blood (Offspring study).**

**S16 Table. RNA-seq, Mild clinical subgroup, Combined 3 litters, DEGs, GOs,Tonsils (Offspring study).**

**S17 Table. RNA-seq, Severe clinical subgroup, DEGs, GOs, Whole blood cells (Offspring study).**

**S18 Table. RNA-seq, Severe clinical subgroup, DEGs, GOs, Tonsils (Offspring study).**

**S19 Table. RNA-seq, HSPCs, DEGs, GOs.**

**S20 Table. DNA methylation array, HSPCs, DEGs, GOs.**

## Notes

### Competing Interest Statement

The authors have declared no competing interest.

