## Supplementary material for "Persistent Japanese encephalitis virus infection in fetuses of an amplifying pig host affects virus genetic heterogeneity and drives transcriptional footprints in offspring": S1_Fig_Maternal_SNVs and iSNVs (Fetal study).docx

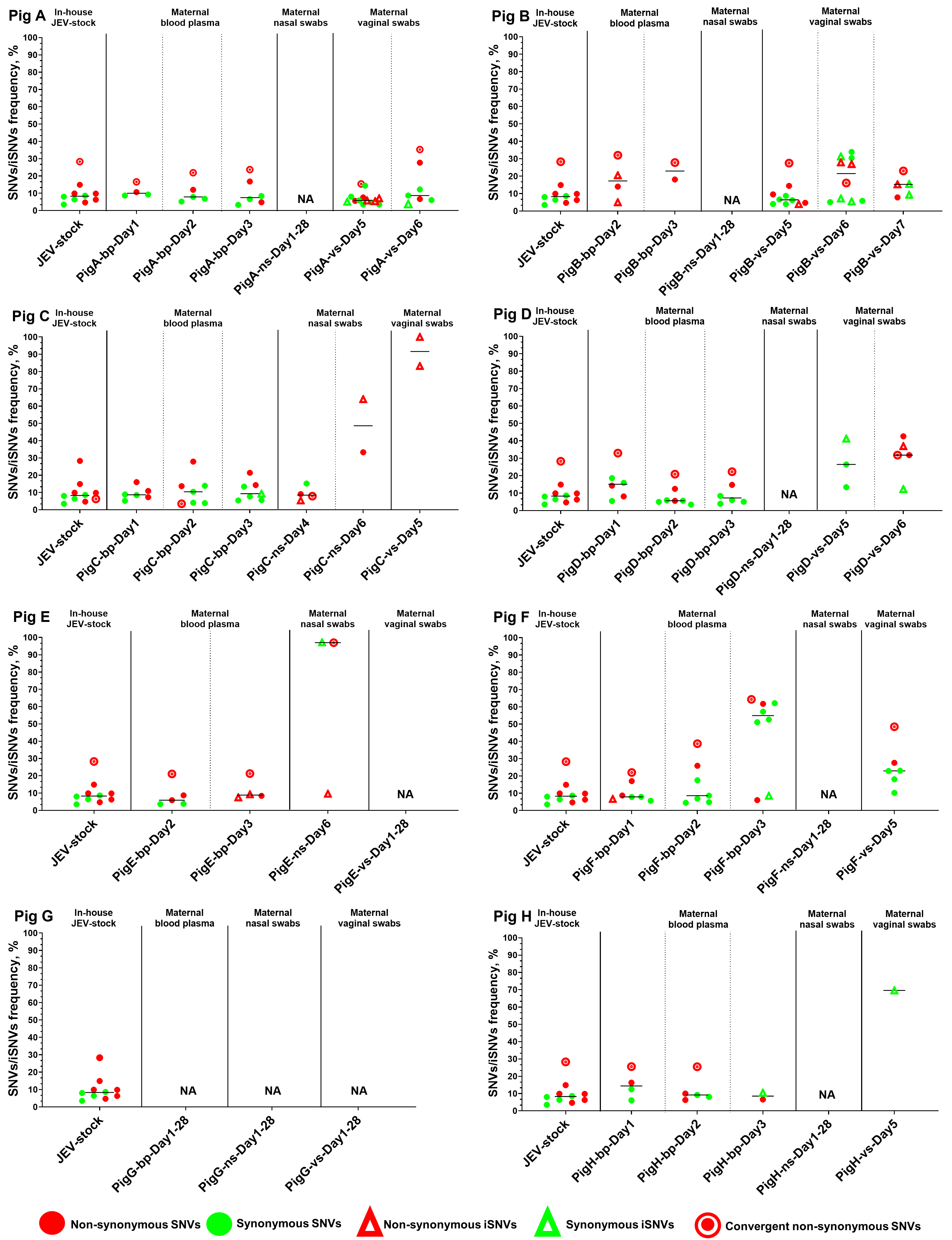

**S1 Fig**. **Single nucleotide variants in samples from adult pregnant and non-pregnant pigs.** **SNVs**: Single nucleotide variants. **iSNVs**: *de novo* emerged intra-host SNVs. **NA**: Library construction for NGS in these samples was not successful. Detailed SNV data is in **S1 Table**. Solid horizontal lines represent median values.
