## Supplementary material for "Persistent Japanese encephalitis virus infection in fetuses of an amplifying pig host affects virus genetic heterogeneity and drives transcriptional footprints in offspring": S1_File_Supplementary_Materials_and_Methods.docx

**RNA extraction, reverse transcription-quantitative polymerase chain reaction (RT-qPCR), and JEV negative-strand-RNA-specific RT-PCR**

We used QIAamp Viral RNA Mini Kit (QIAGEN) according to the manufacturer’s instructions (carrier RNA was not added to avoid interference with NGS) to purify JEV from 140 µl of virus stock, maternal vaginal and nasal swabs (fetal study), maternal (both fetal and offspring study) and fetal (fetal study) blood plasma, and piglet (offspring study) nasal swabs and blood plasma. For nasal and vaginal swabs, swabs (Sterile Puritan HydraFlock Flocked Swabs, #25-3318-H) were inserted into the nose (until slight resistance) or the vagina (approximately 10 cm) and rotated to collect mucosa. Afterward, swabs were placed into tubes containing 500 μl sterile media, the handle was cut one centimeter from the top of the swab, and the tube was stored at -80°C. Before RNA extraction, swab-containing tubes were thawed, shaken for 20 minutes, and spun.

Frozen piglet tissue samples (offspring study—placenta, amniotic membrane, tonsils and brain) were dissected and weighed on analytical balances. One ml of TRI Reagent Solution (Thermo Fisher Scientific) was added to 80–100 mg of tissues before homogenization (5 min at 25 Hz) with RNase-free stainless-steel beads and TissueLyser II (QIAGEN). Then, RNA extraction was performed with PhaseMaker tubes (Thermo Fisher Scientific) and PureLink RNA Mini Kit (Thermo Fisher Scientific) according to the manufacturer’s instructions.

For JEV RNA quantification, we used the previously described probe-based one-step RT-qPCR assay [1-3]. All RT-qPCR reactions were conducted on the StepOne Plus platform (Applied Biosystems) and analyzed using StepOne software version 2.3. The Luna Universal Probe One-Step RT-qPCR Kit (NEB) reaction mixture (20 μL) consisted of 10 μL Luna Universal Probe One-Step Reaction Mix, 1 μL Luna WarmStart RT Enzyme Mix, 1 μL (10 μM) of forward (Universal-JEV-F: 5′-GCCACCCAGGAGGTCCTT-3′) and reverse (universal-JEV-R: 5′- CCCCAAAACCGCAGGAAT-3′) primers, 0.5 μL of the probe (Universal-JEV-Probe: 5′-6FAM-CAAGAGGTG /***ZEN***/ GACGGCC-3′***IABkFQ***), 2.5 μL nuclease-free water and 4 μL of RNA template. The reverse transcription and enzyme activation steps of 10 min at 55°C and 1 min at 95°C were followed by 40 amplification cycles (10 s at 95°C and 60 s at 60°C). A standard curve was used to quantify viral RNA loads as we described [2, 3]. PCR values were normalized to fluid volume or tissue weight, log_10_-transformed, and reported as virus RNA genome copies per ml or g.

Productive infection in tissues was confirmed with JEV negative-strand-RNA-specific RT-PCR [2]: cDNA was synthesized with ProtoScript^®^ II Reverse Transcriptase (New England Biolabs, MA, USA) using 10 μM of the JEV-MinusStr forward primer 5′-GGTCAGAACCACTACTGACAGT-3′. Afterward, cDNA was amplified using 10 μL Luna Universal Probe One-Step Reaction Mix, 3.5 μL nuclease-free water, Universal-JEV-F and Universal-JEV-R (10 μM of each) and Universal-JEV-Probe (10 μM) and 4 μL of cDNA template following the thermal conditions as described above.

In all RNA extraction and PCR assays, we used VERO E6 cell culture media containing JEV as a positive PCR control. As a negative control, we used samples from mock-inoculated pregnant pig and their piglets. Strict precautions were taken to prevent PCR contamination. Aerosol-resistant filter pipette tips and disposable gloves were always used. Kit reagent controls were included in every RNA extraction and PCR run.

**Next-generation sequencing (NGS)**

We used well-established techniques in our laboratory, previously adapted for whole-genome NGS of West Nile virus and Zika virus [2, 4-7]. To generate a consensus whole-genome JEV sequence directly from the virus stock (which was used for inoculation of pigs) we used the *classical non-targeted* NGS protocol for library preparation without PCR pre-amplification. In contrast to samples from JEV infected pigs, the virus stock derived from the cell culture had high viral loads, permitting the use of the non-targeted NGS protocol without advanced PrimalSeq PCR pre-amplification (described below). For non-targeted NGS, RNA was extracted from the viral stocks (140 μl) using the QIAamp Viral RNA Kit (Qiagen) and concentrated via vacuum centrifugation. After vacuum centrifugation, RNA was reverse transcribed into cDNA using the Thermo Scientific Maxima H Minus First Strand cDNA Synthesis Kit (Fisher, MA, USA). The cDNA was further used for the preparation of double-stranded DNA (dsDNA) using E. coli DNA Polymerase I (New England Biolabs), E. coli DNA Ligase (New England Biolabs), RNase H (New England Biolabs), Invitrogen dNTP Mix (10 mM; Thermo Scientific), and NEBNext Second Strand Synthesis (dNTP-free) Reaction Buffer (New England Biolabs). After dsDNA preparation, the samples were purified using NEBNext sample purification beads (New England Biolabs; 2:1 ratio of beads). Afterward, dsDNA was fragmented by NEBNext dsDNA Fragmentase (New England Biolabs) for 20 min at 37°C. DNA concentrations were measured with the Qubit dsDNA HR Assay Kit and Qubit 4.0 Fluorometer (Thermo Fisher Scientific). The purified DNA was used for library construction with the NEBNext Ultra™ II DNA Library Prep Kit for Illumina (#E7645L) and NEBNext Multiplex Oligos for Illumina (96 Index Primers; #E6609S) according to the manufacturer’s instructions. Individual sample libraries were quantified with the Qubit dsDNA HR Assay Kit and Qubit 4.0 Fluorometer. The peak DNA fragment size was identified using the TapeStation HS-DNA 1000 Bioanalysis kit on an Agilent 4150 TapeStation (Agilent). In total, 2 ng of each barcoded library was pooled, quantified (Qubit 4.0), quality-checked (Agilent TapeStation), and converted to moles: Molecular concentration [nM] = Library concentration 84 [ng/μL]/((Average library size x 650)/1,000,000). The pooled library was diluted to 2 nM in 10 mM TE, denatured with 0.1 N NaOH, diluted to 14 pM, and paired-end 300 bp reads were generated with MiSeq Reagent kit v3 (600 cycle output; Illumina) on the MiSeq System (Illumina). To analyze data from non-targeted NGS, the output FASTQ files were quality-trimmed using Trimmomatic 0.39 [8], and the trimmed reads were aligned with BWA to the available reference sequences [GenBank: #EF571853.1]. The sorted and merged BAM files, representing the aligned and assembled NGS reads, were used to calculate NGS coverage and depth using an R script [2, 4-7] (**S3 Table**). Then, we identified the consensus sequence for working JEV stock using Unipro UGENE v46.0 using the default algorithm [9] providing an in-house consensus whole-genome JEV Nakayama reference (**S2 File**).

RNA samples from JEV stock and pig samples (RT-qPCR–positive for JEV) were used for targeted PrimalSeq NGS, which was specifically designed and validated to provide higher sequencing depth, coverage, and sensitivity than classical non-targeted NGS for tissues from natural hosts infected with flaviviruses [10-12]. This protocol enabled, for the first time, near-complete sequencing of flavivirus genomes in tissues from humans and non-human primates [10-12]. The targeted PrimalSeq protocol was applied as we previously described [2, 4-7], using the NEBNext Ultra™ II DNA Library Prep Kit for Illumina (#E7645L) and NEBNext Multiplex Oligos for Illumina (96 Index Primers; #E6609S), and MiSeq Reagent kit v3 (paired-end 600 cycle output; Illumina) on the MiSeq System (Illumina). The PrimalSeq protocol implies the nearly entire virus genome amplified in ~343-363 bp overlapping fragments with multiplexed PCR reactions implying 35 primer pairs (**S1 Table**) [10-12]. JEV cDNA (2 μL) was amplified in two multiplex JEV specific PCR reactions with Primal Scheme primers (**S1 Table**). These two multiplex primer schemes were designed with a web-based tool Primal Scheme [10] using the JEV stock reference sequence (**S2 File**). For NGS data processing and single nucleotide variant (SNVs) calling, we used an open-source software package iVar [10]. The detailed computational protocol used in this study for iVar was previously published by our and other labs [2, 4-7, 10-12]. As the reference sequence for variant calling, we used JEV stock reference sequence (**S2 File**). For the validated best PrimalSeq and iVar practices [10], we analyzed virus genomic regions with a sequencing depth of at least 400×; SNVs with a frequency of at least 0.03 (3%) were considered for analysis; insertions and deletions were not analyzed. In addition, genomic regions amplified with primers that contained nucleotide mismatches within the binding sites were computationally omitted and compensated by overlapping of PCR-amplified fragments. Some pig tissues did not have sufficient coverage and gaps due to low RT-qPCR Ct and JEV loads (**S3 Table**); well-covered regions (at least 400×) in such samples were still analyzed for the presence of SNVs (with a frequency of at least 3%) but these samples were not included into nucleotide diversity and evolutionary selection analysis (described below).

Raw NGS data have been deposited in the Sequence Read Archive under BioProject #PRJNA1467537.

**Japanese encephalitis virus nucleotide diversity analysis**

First, to identify SNVs, we compared PrimalSeq NGS data from JEV stock that was used for pig inoculation and from animal samples to the in-house consensus whole-genome JEV stock reference (**S2 File**). Second, we compared genomic positions and frequencies of SNVs between sequences from JEV stock used for pig inoculation and sequences from maternal, fetal, and piglet samples; tissue-specific SNVs that were not present in the initial JEV stock were considered as *in vivo*-emerged SNVs—iSNVs.

To further confirm that emerged intra-host iSNVs are not virus stock-derived artifacts, we called variants in the JEV stock sequences with the very low 0.01% frequency threshold and compared low-frequency stock-specific SNVs to *in utero*-emerged iSNVs (**S1** and **S7 Tables**). Only iSNVs which did not have corresponding inoculum stock SNVs with a frequency above 0.01% (380 mutations; **S1** and **S7 Tables**) were considered as factual iSNVs.

The percentage of SNVs in virus sequences from each sample was calculated by dividing the number of SNV sites by the total number of sequenced nucleotides. SNV frequencies were also calculated. To quantify the degree of convergent evolution in JEV genome, any iSNV that emerged in more than one animal sample was defined as convergent. And the total convergence was quantified as the percentage of convergent iSNVs from the total number of unique iSNVs.

To further quantify nucleotide diversity metrics (**S1 and S7 Tables**), quality-trimmed paired-end reads were aligned to the indexed JEV stock in-house reference sequence (**S2 File**) using BWA mem v0.7.17. The resulting alignments were converted to BAM format and sorted using SAMtools v1.10 [13] to generate final BAM files for each sample. For samples sequenced in replicates, corresponding BAM files were merged using SAMtools merge, followed by indexing to produce a merged consolidated BAM file per sample. Merged BAM files were used for variant calling without prior normalization for the coverage and depth for each sample. Samples with >50% genome coverage were included in the nucleotide diversity analysis, while a small number of low-coverage samples (four samples) were excluded. Single-nucleotide variants were identified using SAMtools mpileup in combination with VarScan v2.4.4. Mpileup files were generated from each BAM file and processed with VarScan using a minimum coverage threshold of 400×, minimum variant frequency of 3%, p-value cutoff of 0.05, and strand-bias filtering enabled. Variant calls were represented in VCF format for each sample. VCF files were reformatted into SNPGenie-input compatible format using a custom script. Then, SNPGenie v1.2 was used to calculate nucleotide diversity metrics (π, πN, and πS), for each sample by running (snpgenie.pl) from the command line [14].

After the initial analysis, sequencing depth was normalized across samples by random down sampling of BAM files to match the sample with the lowest read count (138,862 reads; mean coverage depth: 3,471) using SAMtools view -s. This ensured normalization of samples and enabled unbiased comparison of nucleotide diversity metrics (π, πN, and πS) across samples using SNPGenie v1.2 [14].

Nucleotide diversity (π) was calculated as the mean number of pairwise nucleotide differences per site across the JEV genome. The mean number of pairwise nonsynonymous differences per nonsynonymous site (πN) and synonymous differences per synonymous site (πS) were also calculated. SNPGenie calculated π, πN, and πS using a statistical method [15] based on that of Nei-Gojobori method [16].

**Reverse genetics to rescue fetal and offspring-specific JEV variants, serial passage to assess stability of acquired mutations, and Sanger sequencing**

We used reverse genetics to synthetically rescue JEV variants containing the fetal-specific C1021T mutation (corresponding to A15V amino acid substitution), and offspring-specific A331G (K79R amino acid substitution) and A6108T (M501L amino acid substitution) mutations. The reasons to select these mutations:

*C1021T (A15V amino acid substitution; fetal-specific)* - This mutation is located in the E protein Domain I β-sheet, which serves as a structural scaffold linking Domains II and III. Residue A15 contributes to the hydrophobic core maintaining E protein integrity during fusion [17, 18]. Substitution to valine likely increases local packing and reduces flexibility, stabilizing the E dimer and enhancing virion stability and infectivity [19]. Reduced E protein flexibility has also been associated with increased neuroinvasiveness, whereas increased flexibility correlates with attenuation [20].

*A331G (K79R amino acid substitution; offspring-specific)* – This mutation lies within the α4 helix (aa 74–96) on the surface of the JEV capsid (C) protein [21]. The C protein consists of four α-helices (α1–α4), of which α4 is involved in RNA binding, nuclear localization, and overall capsid stability [22, 23]. Although not described in JEV, K to R amino acid substitutions in other flavivirus C proteins have been studied. For example, a K101R substitution in the Zika virus C protein markedly increased replication and neurovirulence in mice and induced stronger inflammatory responses compared to the wild-type (WT) virus in neonatal mice [24]. Supporting this, our laboratory showed that an Asian Zika virus acquired a convergent African-lineage K101R mutation in the fetal brain during *in utero* infection in pregnant pigs. The fetal brain–specific variant carrying K101R exhibited higher viral loads in mosquito C6/36 cells and a trend toward increased viral loads in fetal lymph nodes. Notably, the introduced K101R mutation was genetically stable, reaching nearly 100% frequency at 28 days post–*in utero* inoculation in both directly injected and trans-infected fetuses [5].

*A6108T (M501L amino acid substitution; offspring-specific)* – This mutation is identified in offspring samples (**Fig 3F** and **S8 Table**), is located within the C-terminal subdomain III of the JEV NS3 protein, which contributes to the RNA-binding and translocation cleft between helicase domains [25]. Both methionine (M) and leucine (L) are hydrophobic residues; however, L is smaller and less flexible. Thus, the M to L substitution may tighten local packing within the RNA-binding cleft, potentially stabilizing NS3 helicase structure and improving coordination between ATP hydrolysis and RNA translocation [26]. Notably, the M501L substitution has been reported in JEV isolates from bats in China, all belonging to genotype III with high genetic homogeneity (99.4–99.9%) [27] (**Fig 3F**).

The Infectious Subgenomic Amplicons (ISA) reverse genetics method was used as we and others previously described [3-5, 28, 29]. As the reference parental sequence for introduction of fetal-specific (C1021T, amino acid substitution A15V, **S1 Table**) and offspring-specific (A331G, amino acid substitution K79R; and A6108T, amino acid substitution M501L; **S7 Table**) iSNVs, we used the JEV strain SA14-14-2 [GenBank: #MK585066.1]. To rescue JEV variants, we used overlapping (70-80 nt overlap) ISA DNA fragments covering the entire viral genome. We previously published the details of ISA DNA fragments, sequences, and primers for the parental JEV strain SA14-14-2 [3]. The first and last fragments for each virus are flanked with pCMV promoter and HDR/SV40pA sequences. DNA fragments for ISA were ordered through GeneScript (NJ, USA) or Twist Biosciences (CA, USA); correct sequences of ISA DNA fragments were confirmed by NGS.

Overlapping DNA ISA fragments were amplified with Invitrogen Platinum PCR SuperMix High Fidelity (Fisher, MA, USA; #12-532-016), mixed in equimolar concentration to obtain the final 1 µg of DNA for transfection, and transfected into BHK-21 cell monolayers with Lipofectamine 3000 (Fisher, MA, USA; #L3000015) in 640 µl of OptiMEM (Fisher, MA, USA; #11-058-021) for 12 h at +37°C, 5% CO_2_. Afterward, OptiMEM media was removed and replaced with 3 ml of DMEM with 10% of FBS (Fetal Bovine Serum) containing 4×10^5^ Vero cells on top of BHK-21 cells, and plates were incubated for an additional 6 days (passage 0). For passage 1, supernatant from the well with maximum cytopathic effect (CPE) of passage 0 was diluted (1:5000), transferred to a T75 flask with Vero cells, and incubated for 11 days. During passages 0 and 1, cells were monitored for CPE. Generated parental JEV and fetal or offspring-specific JEV variants were passaged 2 times on Vero cells to produce working virus stocks. Cell culture media from the final working stocks were centrifuged (12,000g, 20 min, +4°C), aliquoted, and frozen (-80°C). The absence of mycoplasma contamination in all virus stocks and cell cultures was confirmed using PCR Detection Kit (Millipore Sigma, MA, USA; #MP0035). Genomic sequences in all working stocks and throughout serial passaging were confirmed using Sanger sequencing, as described below.

To define the stability of fetal- and offspring-specific iSNVs, we passaged parental and *in utero*-emerged JEV variants, in Vero cells 5 times. For each subsequent passage, cells were seeded in 6-well plates 24h before virus inoculation and inoculated with 1:100 diluted variants. After 2h at +37°C, the inoculum was discarded, and the cells were covered with fresh culture medium. They were then incubated for 6 days at +37°C and 5% CO_2_. Afterward, the supernatants were collected to inoculate corresponding cells for the next passage. Serial passages for all variants were conducted in two biological replicates.

Viral RNA was extracted from supernatants of parental and variants stocks and at passage 5, as described above. PCR was performed using the Invitrogen SuperScript IV One-Step RT-PCR System (Fisher, MA, USA; #12-594-025) according to the manufacturer’s instructions, with primers spanning sequences encoding parental and modified genomic regions: Fetal iSNV C1021T, amino acid substitution A5V—JEV-SA14-A15V-Fetal-forward: 5′-GTGAATAAAAAAGAGGCTTGGC-3′ and JEV-SA14-A15V-Fetal-reverse: 5′-GCACACATAGCTACTATCAGC-3′). Offspring iSNV A331G, amino acid substitution K79R—JEV-SA14-K79R-piglet-F2: 5′- ATATGCTGAAACGCGGCCTA-3′ and JEV-SA14-K79R-piglet-R2: 5′-CGTCTGCAATGTCCGTGTTG-3’. Offspring iSNV A6108T, amino acid substitution M501L—JEV-SA14-M501L-piglet-forward: 5′- TCATCGACTGTAGAAAGAGCG-3′ and JEV-SA14-M501L-piglet-reverse: 5′- CGGTGTTGTCCTCCAGTATG-3′. PCR products were run on a 1% agarose gel, and DNA was purified and concentrated using DNA Clean & Concentrator-5 (Zymo Research, CA, USA; #D4013) as per the manufacturer’s instructions. DNA was sequenced using Sanger sequencing, and the sequences were aligned with corresponding reference sequences [GenBank: #MK585066.1] and visualized using Unipro UGENE software package.

**RNA-seq and bioinformatics**

RNA samples extracted from piglet whole blood cells and tonsils were used for RNA-seq analysis. RNA from whole blood cells was extracted using Tempus Blood RNA Tubes (Fisher, #4342792) and the Preserved Blood RNA Purification Kit I (Norgen, #SKU43400) according to the manufacturers’ instructions. RNA from tonsils was extracted as described above.

RNA samples were submitted for RNA-Seq at the Donnelly Sequencing Center at the University of Toronto. RNA samples were treated with Turbo DNase. RNA was quantified using Qubit RNA HS (#Q32852, Thermo Fisher Scientific) fluorescent chemistry. RNA Integrity Number (RIN) numbers were assessed by the High Sensitivity RNA ScreenTape (#5067-5579, Agilent Technologies Inc.) and were above 7.  The NEBNext rRNA Depletion Kit v2 (NEB #E7405) was used to deplete ribosomal RNA. In addition, for RNA from whole blood cell samples, we added custom RNA depletion poly A tailed primers (NEBNext Custom RNA Depletion Design Tool v1.0) for Hemoglobin A (5′-GATCTCCGAGGCTCCAGCTTAACGGT-3′; 5′-TCAACGATCAGGAGGTCAGGGTGCAA-3′) and B (5′-AGGGGAACTTAGTGGTACTTGTGGGC-3′; 5′-GGTTCAGAGGAAAAAGGGCTCCTCCT-3′). RNA-seq libraries were prepared from RNA samples (100 ng) using the NEBNext Ultra II RNA Library Prep Kit for Illumina (NEB cat # E7765). The libraries were pooled at equimolar ratios. The final pool was run on an Agilent Bioanalyzer dsDNA High Sensitivity chip and quantified using NEBNext Library Quant Kit for Illumina (NEB #E7630L). The pool at a final concentration of 350 pM was sequenced (paired end 150 bp) on the Illumina NovaSeq6000 platform using a SP flowcell.

Raw FASTQ files were trimmed for adaptor sequences and filtered for low-quality reads using *Trimmomatic*. RNA-seq analysis was conducted as we previously described [2, 4, 30-33]. Paired-end reads were processed using the *kallisto* pseudo-alignment method [34] and quantified by mapping to a transcripts database generated from the pig reference genome assembly [ENSEMBL: #Sscrofa11.1, NCBI RefSeq assembly: #GCA_000003025.6]. The count table for RNA-seq data was assembled using the *tximport::tximport* function in R. After importing data to the R environment, we removed data for genes with low expression using the *edgeR::filterByExpr* function. Normalization was then performed using the *edgeR::calcNormFactors* function, and the *limma::voom* function was used to convert the data into a normal distribution. We calculated differential expressions using the *limma::lmFit* function with empirical Bayes moderation via the *limma::eBayes* function. Finally, we performed a Gene Set Enrichment Analysis (GSEA) using the *limma::camera* function with Gene Ontology (GO) annotation for *Sus scrofa* obtained from g:Profiler [35, 36]. We compared gene expression directly between Mild (or Severe) clinical subgroup and the Control group. Direction of analysis for RNA-seq was Mild clinical subgroup/Control group or Severe clinical subgroup/Control group. Differentially expressed genes with FDR < 0.05 and log_2_FC ≥ 0.5 were considered significant. GO Biological processes with FDR < 0.25 were considered significant. RNA-seq data are in result figures or in **S9-18 Tables**. Raw sequencing data were deposited to NCBI BioProject under accession number #PRJNA1467734.

**Experiments with CD34^+^ hematopoietic stem and progenitor cells (HSPCs)**

The work was conducted with human HSPCs because, in contrast to pig cells, this population is well characterized and there are readily available reagents and established protocols. For work with human samples, we followed the University of Saskatchewan Human Research Ethics Policy with approval #16-135 from a Biomedical Research Ethics Board (Bio-REB). Eleven pregnant women recruited (27^th^ August 2020–25^th^ November 2020) into the study were free from hepatitis B, C, and human immunodeficiency virus. All participants provided written informed consent. Ten to twenty ml of umbilical cord blood was collected from eleven healthy full-term deliveries with six female and five male newborns. Blood was collected immediately after delivery from the umbilical cord vein using BD Vacutainer and EDTA-containing Blood Collection Tubes. Blood samples were stored on ice packs and delivered to the lab within an hour after sampling. Blood was diluted 1:2 in Dulbecco’s phosphate-buffered saline (DPBS; Gibco) and peripheral blood mononuclear cells were isolated by centrifugation (400g, 30 min, +20 °C, using slow acceleration and brake settings) over Lymphocyte Separation Medium (Corning) followed by three washes in DPBS. To isolate CD34+ cells, we used magnetic-activated cell sorting according to the manufacturer’s instructions for Miltenyi Biotec human CD34 MicroBead Kit and LS Columns with an elution buffer (commercial DPBS + 2 mM EDTA + 0.5% BSA). The enriched CD34+ hematopoietic progenitor stem cell (HSPC) population was washed (300g, 5 min, +4°C) with StemSpan™ SFEM medium (STEMCELL Technologies Inc.) and counted. On average, 2-5×10^5^ CD34+ HSPCs were recovered from umbilical cord blood samples.

We tested how exposure to IFN-α affects the differentiation of fetal CD34+ HSPCs into granulocyte-macrophage progenitors (GMPs) and molecular responses in HSPCs-derived GMPs. We also tested LPS-induced cytokine response in HSPCs-derived GMPs. The experimental setup is represented in **S3 Fig**.

After magnetic-activated cell sorting, fetal CD34+ HSPCs were exposed to human recombinant IFN-α (PHC4014, Thermo Fisher Scientific) in StemSpan™ SFEM medium supplemented with 1× StemSpan™ CD34+ Expansion Supplement (STEMCELL Technologies Inc.), 1× P/S (Penicillin/Streptomycin; Gibco), 50 μg/ml Gentamycin (Thermofisher), and 175 nm UM171 (STEMCELL Technologies Inc.) at +37°C. For exposure, HSPCs were plated into the 96-well plates (5×10^4^ cells per well) in the presence of 1000 U/ml IFN-α. Control cells were plated without IFN-α. Exposed and Control cells were harvested after 24 hours by gentle pipetting, washed two times (300g, 10 min, 4°C) in Iscove's Modified Dulbecco's Medium (IMDM; Thermo Fisher Scientific) with 2% FBS, resuspended in the same media, and counted. Afterward, for differentiation, IFN-α-treated (for 24 h) and Control CD34+ cells were subjected to the colony-forming unit assay.

In the colony-forming unit assay, IFN-α-treated (for 24 h) CD34+ HSPCs were differentiated for 10 days with or without IFN-α, resulting in two differentiation conditions—IFN-α-short-treated (24 h with IFN-α and then 10 days with no IFN-α), and IFN-α-long-treated (24 h with IFN-α and 10 days with IFN-α) **S3 Fig**. Control CD34+ HSPCs were not exposed to IFN-α at any experimental stage. The colony-forming unit assay is the classical assay to measure the proliferation and differentiation of hematopoietic stem/progenitor cells in mice, pigs, and humans [33, 37-39]. For this, 800 µl of CD34+ HSPC suspension containing 18,000 cells was first mixed with 8 ml of MethoCult™ H4534 Classic Without EPO media (STEMCELL Technologies Inc.), and then aliquoted with a sterile 18-gauge needle and 1 mL syringe into 6-well plates (1.1 ml per well; six-well replicates in SmartDish™ 6-well plates, STEMCELL Technologies Inc.) that resulted in 2,250 cells per well. For the IFN-α-long-treated condition, an initial 800 µl cell suspension was supplemented with human recombinant IFN-α resulting in the final IFN-α concentration of 1000 U/ml after mixing with MethoCult™ media. The concentration of IFN-α was selected based on previous studies [38, 39]. For IFN-α-short-treated and Control conditions, IFN-α was not added in the colony-forming unit assay. After 10 days, granulocyte-macrophage colonies were morphologically identified with an inverted light microscope according to the StemCell Technologies Atlas of Human Hematopoietic Colonies (STEMCELL Technologies Inc.). The total number of granulocyte-macrophage colony-forming units (CFU-GM) in all six wells was counted using STEMgrid™-6 (STEMCELL Technologies Inc.). After counting CFU-GM, colonies were harvested and disrupted into single-cell suspension, and the total number of granulocyte-macrophage progenitors (GMPs) was calculated.

To harvest CFU-GM, plates were chilled at 4°C for 1 hour, and MethoCult™ media with cells from 6 well replicates were resuspended in a cold RPMI medium (2 ml per well) containing 1× P/S, 1 mM sodium pyruvate and 50 μg/ml Gentamycin. Afterward, cells were washed two times (400g, 15 min, 4°C), resuspended in RPMI media, and counted. The portion of GMPs—10^6^ cells, was used to concurrently extract and separate RNA (for RNA-seq) and DNA (for whole-genome methylation assay) with RNA/DNA Purification Kit (Norgen Biotek) according to the manufacturer’s instructions. Another portion of granulocyte-macrophage progenitors—1.5×10^6^ cells, was used for activation with LPS. For this, cells were plated in triplicate (2×10^5^) in 96-well plates with RPMI medium containing 1× P/S, 1 mM sodium pyruvate, 50 μg/ml Gentamycin and 1 μg /ml LPS (Sigma-Aldrich). Control cells were plated in triplicate in the same media but with no LPS. After 24 hours, supernatants from LPS-treated and Control cells were collected, centrifuged (2,000g, 5 min, 4°C), and stored at -80°C until analysis. To analyze LPS-induced activation, we quantified IL-1*β* in supernatants using an enzyme-linked immunosorbent assay (ELISA) kit for human IL-1*β* (R&D).

RNA was isolated from granulocyte-macrophage progenitors as described above and was assessed on a bioanalyzer; all samples had RNA Integrity Number RIN values above 8.5. DNA from samples was removed with TURBO DNA-free™ Kit (Thermo Fisher Scientific). Then, mRNA with intact poly(A) tails were enriched with NEBNext Poly(A) mRNA Magnetic Isolation Module (New England Biolabs) and used for library constructions with NEBNext® Ultra™ II Directional RNA Library Prep Kit for Illumina and NEBNext Multiplex Oligos for Illumina (96 Unique Dual Index Primer Pairs; New England Biolabs).

Libraries were sequenced on the NovaSeq as paired-end reads using the NovaSeq 6000 S1 Reagent Kit v1.5 100 cycles (Illumina). FASTQ files were trimmed for adaptor sequences and filtered for low-quality reads using *Trimmomatic*. On average, 15.1 million reads per sample were generated. RNA-seq analysis was performed as we previously described [2, 4, 30-33]. Briefly, a complete transcriptome database was generated from ENSEMBL *Homo sapiens* GRCh38.p13 (GCA_000001405.28). Sequencing data were mapped and quantified using *kallisto* [34]. Then counts were analyzed using *Bioconductor* packages *tximport*, *edgeR*, and *limma*. The *voom* function from the *limma* package was used for differential expression analysis. Gene set enrichment analysis was performed with *camera* function in *limma* using the GMT file (v7.5.1) containing symbols of gene sets derived from the Gene Ontology Biological Process Ontology of the Gene Set Enrichment Analysis (GSEA) Molecular Signatures Database (MSigDB) (**S19 Table**).

The set enrichment results from *camera* were graphed in *Cytoscape* using the *EnrichmentMap* plugin [32, 40]. All networks were generated using a Jaccard + Overlap with a cutoff of 0.375 and a Combined Constant of 0.5. Sub-networks were discovered using GLay cluster and annotated using the WordCloud plugin of the top 4 words with a bonus of 8 for word co-occurrence. An accession number for RNA-seq data is #PRJNA836933 in NCBI BioProject.

Granulocyte-macrophage progenitors from four patients were analyzed in whole-genome DNA methylation array. Quantity and quality of genomic DNA were assessed with Qubit and NanoDrop 8000 (Thermo Fisher Scientific). We used 750 ng of DNA for bisulfite conversion with the EpiTect Bisulfite Kit (QIAGEN) and methylation microarray with the Infinium MethylationEPIC BeadChip Kit (Illumina); scanning was done by an Illumina iScan. The Infinium MethylationEPIC BeadChip Kit covers over 850,000 methylation sites quantitatively across the genome at single-nucleotide resolution (Illumina).

Methylation data processing, identification of differentially methylated positions, and epigenetic regulation of gene expression were performed as we previously described [40] with modifications. Briefly, array data were extracted from image files and processed for normalization and background correction using Illumina GenomeStudio v2011.1 (MethylationEPIC v-1-0 B3 annotation file was used). Text tables containing sample information and probe signals were processed using the methylumi_2.42.0 package in R v4.2 and annotated to the genome using the IlluminaHumanMethylationEPICanno.ilm10b2.hg19 library in R. We compared whole-genome methylation between Control and long-term IFN-α treatment conditions in granulocyte-macrophage progenitors from four patients to identify differentially methylated positions, using the *limma* library [40, 41]. All differential methylation analysis was performed using M values calculated by beta/(1-beta) as this is known to improve the statistical calculation of differential methylation [42, 43]. Differential methylation was calculated using the limma v3.52.2 package in R accepting FDR adjusted *P*-values < 0.05 and log_2_FC > 1 as significant (**S20 Table**). Gene set overrepresentation analysis was performed using Human Mine (www.humanmine.org) on the top demethylated genes (FDR corrected *P*-value < 0.05). Gene Ontology terms with an FDR corrected *P*-value < 0.05 were held as significant (**S20 Table**). Raw data from the methylation microarray with the Infinium MethylationEPIC BeadChip Kit (Illumina) and Illumina iScan were deposited in DRYAD: <https://doi.org/10.5061/dryad.v6wwpzhbq>.
