## Supplementary material for "Persistent Japanese encephalitis virus infection in fetuses of an amplifying pig host affects virus genetic heterogeneity and drives transcriptional footprints in offspring": S2_Fig_Fetal_SNVs and iSNVs (Fetal study).docx

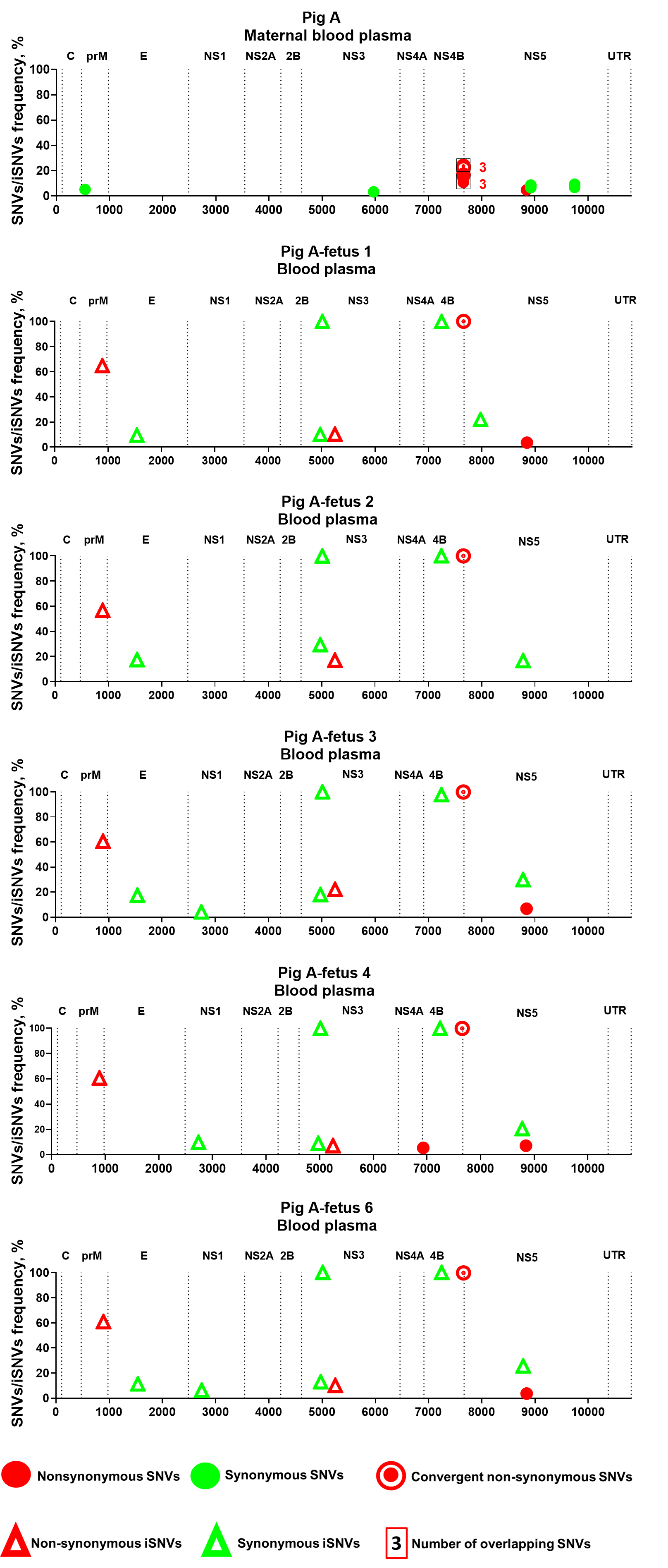
**S2 Fig**

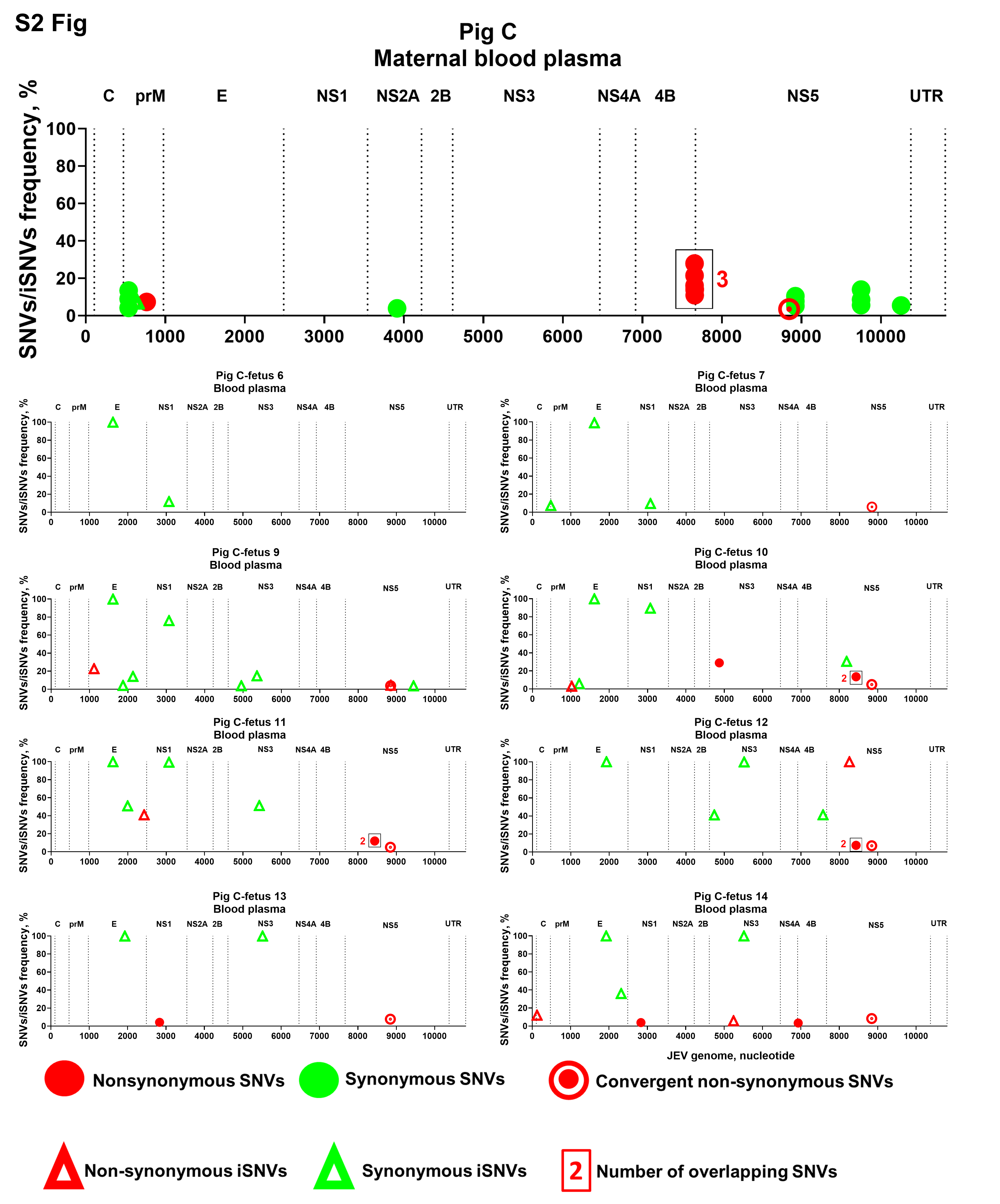

**S2 Fig. Nucleotide position of SNVs and iSNVs within the JEV genome.** Pattern of JEV SNVs and iSNVs in the maternal and fetal blood from pregnant **Pig A** and **Pig C**, and their fetuses. **C**: JEV capsid protein. **prM**: Precursor membrane protein. **E**: Envelope protein. **NS1**: Nonstructural protein 1. **NS2A**: Nonstructural protein 2A. **NS2B**: Nonstructural protein 2B. **NS3**: Nonstructural protein 3. **NS4B**: Nonstructural protein 4B. **NS5**: nonstructural protein 5. **UTR**: untranslated region. Raw data is shown in **S1 Table**.
