## Supplementary figures and images for "Persistent Japanese encephalitis virus infection in fetuses of an amplifying pig host affects virus genetic heterogeneity and drives transcriptional footprints in offspring"

### S3 Fig. The HSPC experimental setup.jpg

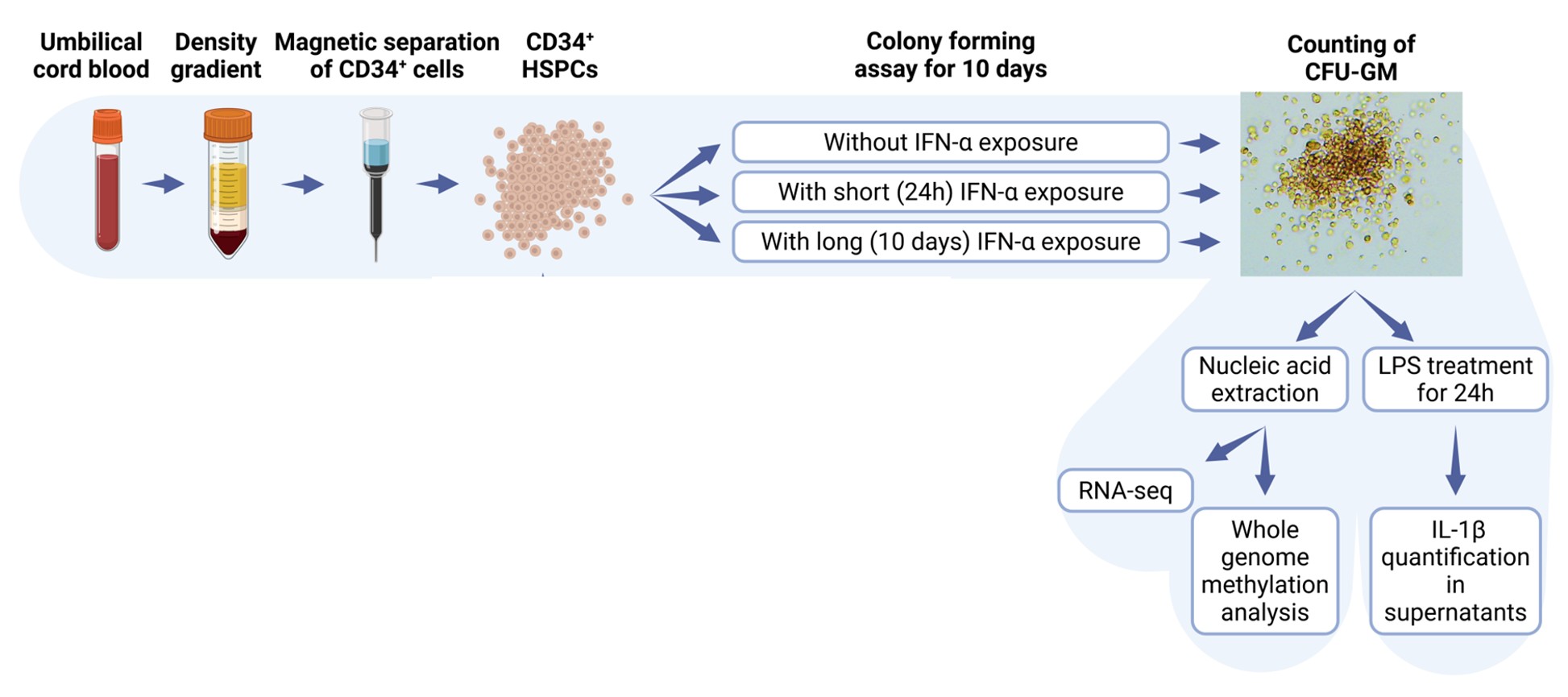
